# Plasma Glycomic Markers of Accelerated Biological Aging During Chronic HIV Infection

**DOI:** 10.1101/2023.08.09.551369

**Authors:** Leila B Giron, Qin Liu, Opeyemi S Adeniji, Xiangfan Yin, Toshitha Kannan, Jianyi Ding, David Y. Lu, Susan Langan, Jinbing Zhang, Joao L. L. C. Azevedo, Shuk Hang Li, Sergei Shalygin, Parastoo Azadi, David B Hanna, Igho Ofotokun, Jason Lazar, Margaret A. Fischl, Sabina Haberlen, Bernard Macatangay, Adaora A. Adimora, Beth D. Jamieson, Charles Rinaldo, Daniel Merenstein, Nadia R. Roan, Olaf Kutsch, Stephen Gange, Steven Wolinsky, Mallory Witt, Wendy S. Post, Andrew Kossenkov, Alan Landay, Ian Frank, Phyllis C. Tien, Robert Gross, Todd T. Brown, Mohamed Abdel-Mohsen

**Author notes:** Corresponding author: Mohamed Abdel-Mohsen, Ph.D. Associate Professor, Vaccine and Immunotherapy Center, The Wistar Institute. 3601 Spruce Street, Philadelphia, PA 19104.

## Abstract

People with HIV (PWH) experience an increased vulnerability to premature aging and inflammation-associated comorbidities, even when HIV replication is suppressed by antiretroviral therapy (ART). However, the factors that contribute to or are associated with this vulnerability remain uncertain. In the general population, alterations in the glycomes of circulating IgGs trigger inflammation and precede the onset of aging-associated diseases. Here, we investigate the IgG glycomes of cross-sectional and longitudinal samples from 1,216 women and men, both living with virally suppressed HIV and those without HIV. Our glycan-based machine learning models indicate that living with chronic HIV significantly accelerates the accumulation of pro-aging-associated glycomic alterations. Consistently, PWH exhibit heightened expression of senescence-associated glycan-degrading enzymes compared to their controls. These glycomic alterations correlate with elevated markers of inflammatory aging and the severity of comorbidities, potentially preceding the development of such comorbidities. Mechanistically, HIV-specific antibodies glycoengineered with these alterations exhibit reduced anti-HIV IgG-mediated innate immune functions. These findings hold significant potential for the development of glycomic-based biomarkers and tools to identify and prevent premature aging and comorbidities in people living with chronic viral infections.

## INTRODUCTION

Even with long-term suppressive antiretroviral therapy (ART), people with chronic HIV infection (PWH) prematurely experience a high incidence of aging-associated diseases, including cardiovascular disease (CVD), cancers, and neurocognitive disorders.^1^ There are considerable gaps in our understanding of the pathophysiological mechanisms driving the development of such comorbidities in PWH; however, many of these comorbidities are linked to a chronic inflammatory state called inflammaging, commonly observed in elderly individuals.^2–4^ The precise mechanisms driving inflammaging in PWH are not fully understood, but they may involve ongoing HIV production, cytomegalovirus (CMV) infection, loss of regulatory T cells, microbial translocation, and other undetermined host and viral factors.^5^ Comprehensive understanding of the factors associated with inflammaging in PWH can facilitate the development of biomarkers to predict the occurrence or severity of inflammaging-associated comorbidities in PWH, and may aid in the development of tools to prevent the onset of these comorbidities.

Aberrant host glycosylation has recently emerged as a key driver of chronic inflammation and accelerated biological aging in the general population.^6–9^ Within the circulating glycome, the glycans on antibodies are especially critical as these are linked to systemic and chronic inflammatory responses. Specifically, the glycans on the Fc domain of circulating immunoglobulins G (IgGs) play a crucial role in regulating antibody non-neutralizing functions, including antibody-dependent cell-mediated cytotoxicity (ADCC), antibody-dependent cellular phagocytosis (ADCP), complement-dependent cytotoxicity (ADCD), and various pro- and anti-inflammatory activities.^10,11^ This connection between IgG glycosylation and systemic inflammatory responses has particularly emphasized the impact of antibody galactosylation and sialylation in promoting strong anti-inflammatory effects.^12–14^ For instance, sialic acid-containing *N*-linked glycans were shown to mediate the potent anti-inflammatory properties of intravenous immunoglobulin (IVIG).^13,15–17^ Conversely, a decrease in IgG sialylation, known as hypo-sialylation, enhances the pro-inflammatory function of IgGs.^12–14^

Various characteristics of IgG glycosylation have also been strongly associated with both chronological and biological aging in various large-scale studies conducted in the general population. For example, specific glycomic characteristics of IgG, particularly the loss of galactose, known as agalactosylation, can better predict chronological and biological age than traditional markers such as telomere length.^18,19^ Glycomic traits have also been found to be significantly altered in individuals with age-related illnesses, including inflammatory bowel disease, systemic lupus erythematosus, CVD, cancer, and diabetes.^20–25^ Whether IgG glycosylation drives, or is simply a biomarker of aging, aging-related diseases, and aging-related comorbidities is still a subject of debate. However, changes in IgG glycosylation have been observed years before the onset of some diseases, indicating a potential causative role.^26,27^

Despite the growing body of evidence suggesting a connection between an altered circulating IgG glycome and accelerated biological aging, it remains unclear whether chronic HIV infection accelerates the pace of aging-associated IgG glycomic alterations. Our group previously showed that IgGs of PWH had lower levels of circulating glycans suggested to be anti-inflammatory and anti-aging (i.e., sialylated and galactosylated glycans) than did IgGs of HIV-negative individuals.^28^ However, in that prior study, the sample size was small, and the groups were not matched for demographic factors such as age, sex, and ethnicity. In addition, that study did not examine the potential upstream mechanisms of these alterations or the potential downstream consequences of them. In this current study, we aim to address these limitations.

## RESULTS

### Long-term, ART-suppressed HIV infection is associated with sex-dependent IgG glycomic alterations

We first sought to examine whether well-suppressed HIV infection impacts the IgG glycomes. To isolate the influence of ART-suppressed HIV infection on the IgG glycomes, while minimizing confounding factors and selection biases, we selected cross-sectional samples from PWH on suppressive ART and HIV-negative counterparts enrolled in the MACS/WIHS Combined Cohort Study (MWCCS). The key feature of this cohort is the comparable recruitment, data collection procedures, and long-term retention of PWH and well-matched HIV-negative controls.^29–32^ As it is documented that sex impacts IgG glycosylation,^33^ we separated our analysis by sex assigned at birth, specifically biologically-born women (hereafter referred to as women) and biologically-born men (hereafter referred to as men). We analyzed the IgG glycomes of 254 women with HIV (WWH) on suppressive ART for at least five years with undetectable HIV viral load, weight less than 300 pounds, and with a median CD4 T cell count of 726 cells/mm^3^. We matched these samples with samples from 235 HIV-negative women matched to the WWH in terms of age, race, and body mass index (BMI) (**Table 1**, **Figure 1A**). To ensure a wide age range for analysis, we aimed to have a similar number of individuals in each of the following age categories: ≤45, 46-50, 51-55, 56-60, 60-65, and >65 years (**Table 1**, **Figure 1A**). Using the same criteria, we also analyzed samples from 243 men with HIV (MWH) who were suppressed on ART with a median CD4 T cell count of 698 cells/mm^3^ and 253 HIV-negative matched controls (**Table 1**, **Figure 1A**).

**Table 1.**
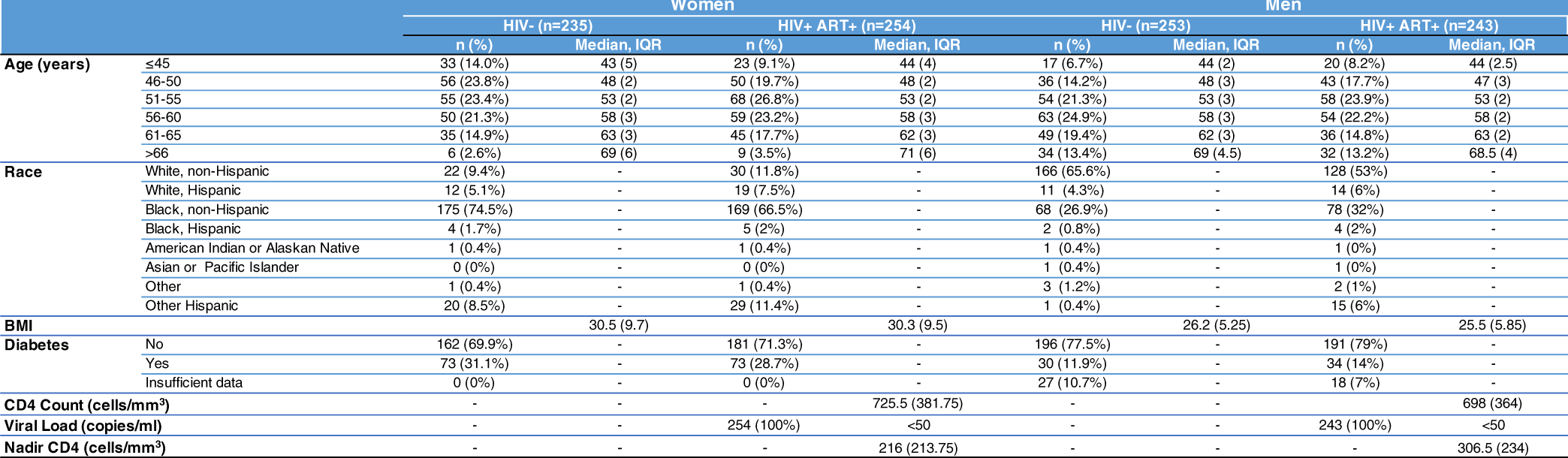
The characteristics of participants in studies depicted in Figures 1-4 and 8.

**Figure 1.**
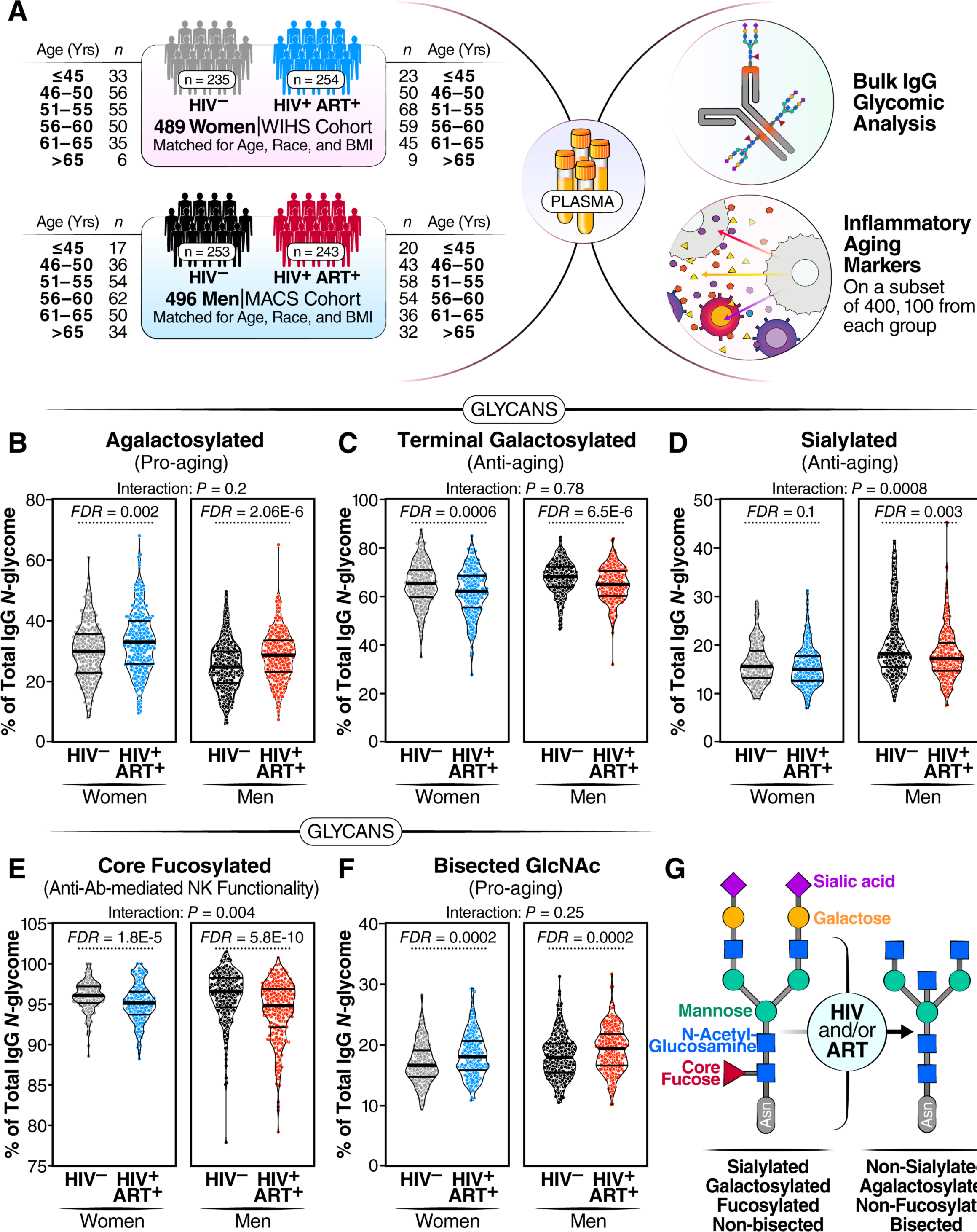
Long-term ART-suppressed HIV infection is associated with sex-dependent IgG glycomic alterations. **(A)** Overview of the main study design. **(B-F)** Violin plots displaying the levels of IgG glycan groups in WWH and MWH undergoing long-term ART compared to their HIV-negative counterparts. The median and interquartile range (IQR) are shown. Unpaired t-tests and false discovery rates (FDRs) were calculated to account for multiple tests over studied markers. Interaction *P* values were calculated using multivariable models, adjusting for age, ethnicity, and BMI. **(G)** A schematic summary illustrating the IgG glycomic alterations associated with HIV infection and/or ART treatment.

We identified 21 individual *N*-glycan structures within the bulk IgG glycomes of these samples (**Supplementary Figure 1A**). These glycan structures were grouped into five IgG glycomic groups, depending on the presence or absence of four key monosaccharides: sialic acid (sialylated), galactose (agalactosylated, and terminal galactosylated), fucose (fucosylated), and bisecting GlcNAc (bisected) (**Supplementary Figure 1B**).

We first assessed whether ART-suppressed HIV infection is associated with alterations in the levels of any of these IgG glycan groups, compared to controls, and whether the degree of alteration differed between WWH and MWH. We found that ART-suppressed HIV infection was associated with higher levels of agalactosylated glycans (pro-aging) in both men and women (FDR<0.05; **Figure 1B**). To examine whether this induction of agalactosylated glycans differed between women and men, we needed to account for potential confounding factors such as ethnicity and BMI, which differed between the women and men (**Table 1**). We employed a multi-variable model that adjusted for age, ethnicity, and BMI. The interaction comparison revealed no significant differences in the association between HIV/ART and IgG agalactosylated glycans between women and men (**Figure 1B**). These findings suggest that HIV and/or ART induce similar levels of agalactosylated glycans (pro-aging) in WWH and MWH compared to HIV-negative counterparts. Using a similar analysis approach, we examined the effects of HIV/ART on the other IgG glycan groups. We found that HIV/ART was associated with: 1) a reduction of terminally galactosylated glycans (anti-aging) in both men and women (**Figure 1C**), 2) a reduction of sialylated glycans (anti-inflammatory) in men but not women (**Figure 1D**), 3) a reduction of core fucosylated glycans (which are involved in the modulation of IgG-mediated ADCC), in both men and women, but more profoundly in men than women (**Figure 1E**), and 4) an induction of the bisected GlcNAc glycans (pro-aging) in both men and women (**Figure 1F**), compared to HIV-negative counterparts. A detailed comparison of the 21 individual glycan structures (**Table 2**) supported the group analysis that HIV and/or ART alter the levels of several of these glycan structures.

**Table 2.**
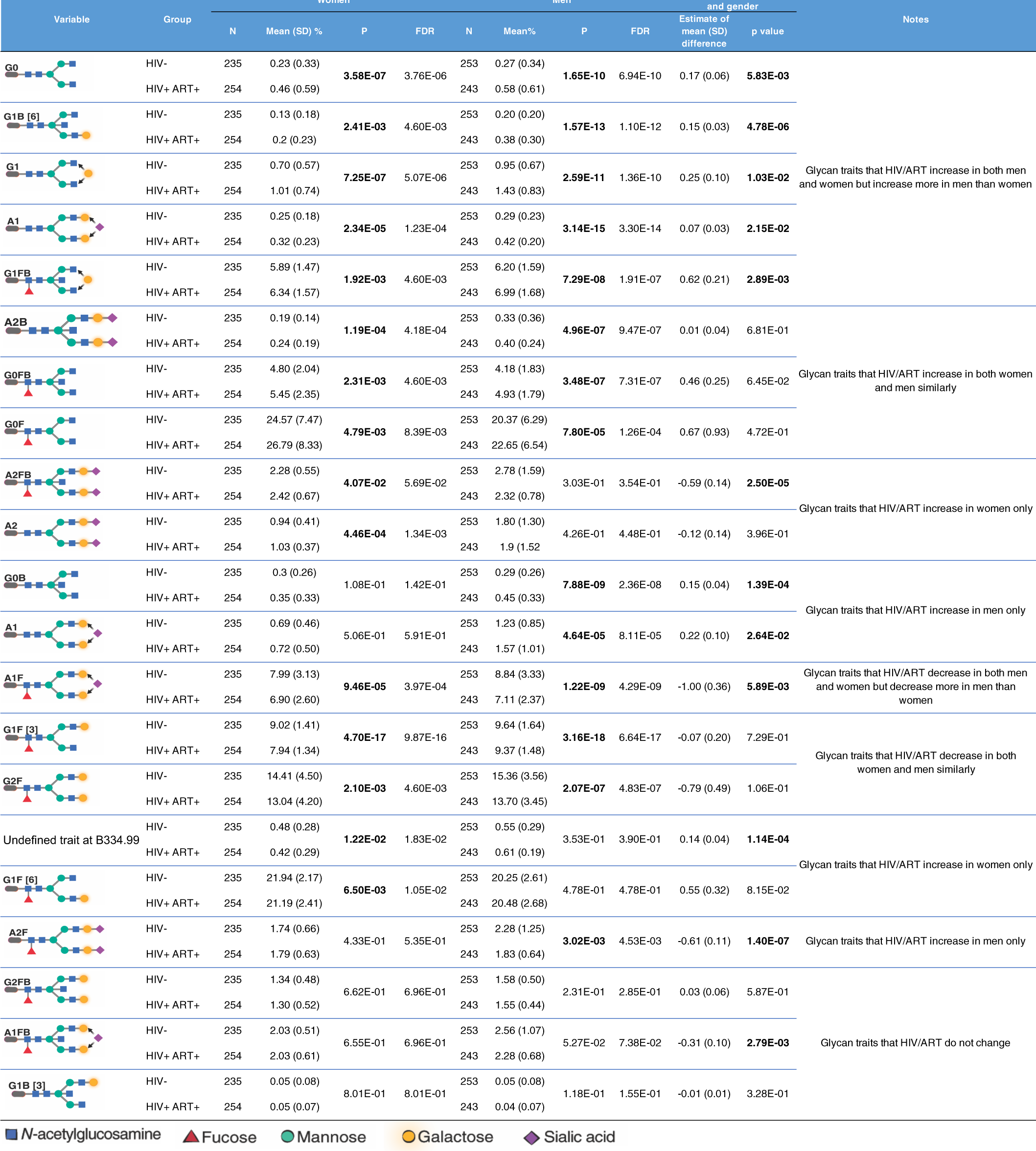
IgG glycomic traits modulated by living with HIV and/or ART.

Because previous studies in the general population indicated that menopause and sex hormones have an impact on IgG glycosylation,^33–36^ we performed a secondary analysis of the women samples based on self-reported menopause status: pre-menopause (n=60 HIV-negative and n=35 WWH), peri-menopause (n=21 HIV-negative and n=20 WWH), or post-menopause (n=152 HIV-negative and n=198 WWH). We compared the IgG glycomic groups (**Supplementary Figure 2**) or individual glycan traits (**Supplementary Table 1**) among these menopause groups. Our findings confirmed that menopause and/or age are associated with a reduction in glycan structures that are considered anti-aging and anti-inflammatory (galactosylated and sialylated) and an increase in glycan structures that are considered pro-aging and pro-inflammatory (agalactosylated and bisected), in both WWH and their HIV-negative counterparts.

Together, these findings suggest that chronic virally-suppressed HIV infection is associated with distinct and likely detrimental IgG glycomic alterations. These alterations include an accumulation of pro-aging glycans (agalactosylated and bisected GlcNAc), a decline in anti-aging glycans (terminally galactosylated and sialylated), and a decrease in glycans involved in the modulation of ADCC (fucosylated), with variations observed based on sex (**Figure 1G**).

### HIV/ART-mediated IgG glycomic alterations correlate with higher markers of inflammatory aging

IgG glycomic alterations have been linked to inflammatory activities.^12–17,37–39^ Considering that ART-suppressed HIV infection significantly impacts IgG glycomes, our next objective was to investigate whether the changes in IgG glycomes associated with HIV/ART are correlated with inflammation. To achieve this, we selected 400 individuals, with 100 from each of the four groups in Figure 1A. These were matched for age and ethnicity between groups of women and men. We measured the levels of 22 inflammatory markers known to play crucial roles in HIV pathogenesis (such as sCD14, sCD163, IL-6),^5^ as well as markers associated with inflammatory aging.^40^ Among the markers associated with inflammatory aging, we included three markers (CXCL9, Eotaxin, and Leptin), which had been recently incorporated into an inflammatory aging clock capable of predicting accelerated biological aging in the general population.^40^ We found that, in general, PWH exhibit higher levels of plasma inflammatory markers in a sex-dependent manner. For instance, IP-10 and sCD14 were elevated in women and men living with HIV compared to their controls. CXCL9 was higher in WWH but not in MWH compared to their respective controls. Additionally, levels of sCD163, IL-4, IL-5, IL12p70, MIP-1α, and MCP-2 were elevated in MWH but not in WWH compared to their controls (**Figure 2A-I**). We also investigated if these markers differed among women based on menopause status. Menopause was associated with increased levels of CXCL9 and Eotaxin, and the association of HIV infection on CXCL9 and IP-10 was more pronounced in post-menopausal women compared to pre-menopausal women (**Supplementary Figure 3**).

**Figure 2.**
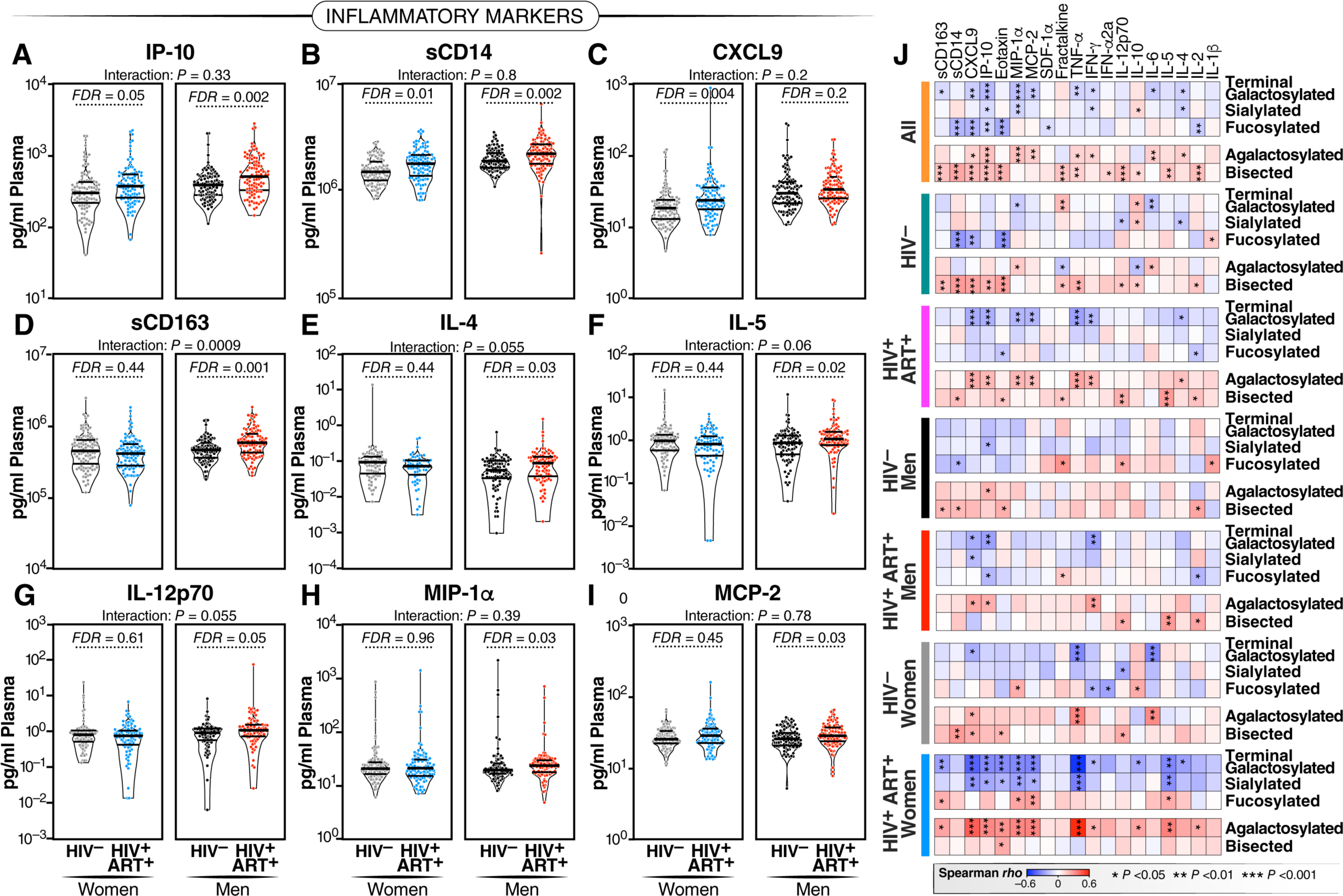
HIV/ART-mediated IgG glycomic alterations correlate with higher markers of inflammatory aging. **(A-I)** Violin plots illustrating the levels of various plasma markers of inflammation in WWH and MWH undergoing long-term suppressive ART compared to their HIV-negative counterparts. Median and IQR are displayed. Unpaired t-tests and FDRs were employed to address multiple comparisons. Interaction *P* values were computed using multivariable models, adjusting for age, ethnicity, and BMI. **(J)** Spearman’s rank correlation heatmaps revealing the associations between IgG glycan groups (rows) and several plasma markers of inflammation (columns) in different groups of individuals (all samples combined, HIV-negative individuals, PWH, MWH, HIV-negative men, WWH, HIV-negative women). Positive correlations are depicted in red, while negative correlations are shown in blue. * P<0.05, ** P<0.01, and *** P<0.001.

Given that both IgG glycans and inflammatory aging markers differed by HIV serostatus, we then examined correlations. As depicted in **Figure 2J**, we found negative correlations between the glycans that are depleted during chronic HIV infection (sialylated, galactosylated, and fucosylated glycans; Figure 1G) and multiple markers of inflammatory aging. Conversely, we found positive correlations between the glycans that are enriched during chronic HIV infection (agalactosylated and bisected GlcNAc glycans; Figure 1G) and markers of inflammatory aging. These correlations were evident when considering the entire cohort, HIV-negative controls, PWH, MWH, and WWH. These data suggest that HIV-associated IgG glycomic alterations are linked to higher inflammation, aligning with their known functions.

### The links between IgG glycans and chronological age are altered in PWH

Previous studies in the general population indicated that specific IgG glycans correlate with chronological age.^18^ However, given that ART-suppressed HIV infection is associated with changes in the IgG glycome, we next sought to determine whether the HIV/ART-mediated alteration of IgG glycomes altered the relationship between IgG glycomes and chronological age. We identified three distinct patterns. First, several glycan structures exhibit a positive correlation with age in both PWH and their HIV-negative counterparts. However, the slope of these correlations is significantly different in PWH, suggesting that HIV either accelerates or decelerates the accumulation of these glycan structures over time. For instance, the agalactosylated G0 glycan trait (**Figure 3A**) and the bisected GlcNAc grouped glycans (**Figure 3B**) both positively correlated with age in PWH and HIV-negative individuals, but the slope of these correlations differs significantly between the two groups (*P*=0.0075 and *P*=0.059, respectively). Second, specific glycan traits, such as G2FB (**Figure 3C**) or A1 (**Figure 3D**), do not correlate with age in one of the two populations but do correlate in the other population (e.g., G2FB only in PWH, or A1 only in HIV-individuals). Third, specific glycans such as G2F (**Figure 3E**) show a consistent correlation with age in both PWH and their HIV-negative counterparts. The complete dataset can be found in **Supplementary Table 2**.

**Figure 3.**
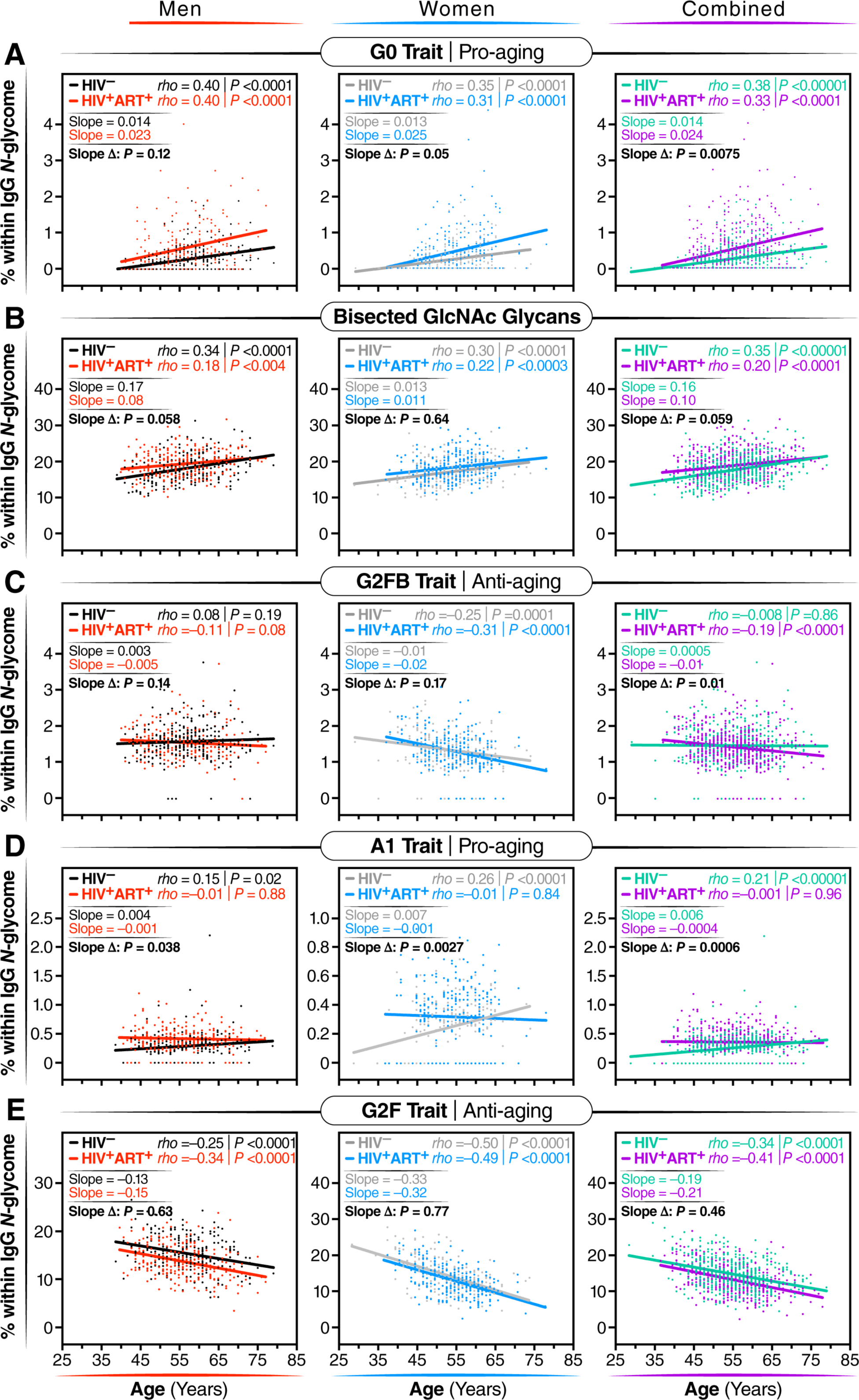
The associations between IgG glycans and chronological age are altered in individuals with long-term ART-suppressed HIV infection. Spearman’s rank correlations between chronological age (x-axis) and the levels of the following glycan traits (y-axis): **(A)** G0 glycan trait, **(B)** bisected GlcNAc glycans, **(C)** G2FB glycan trait, **(D)** A1 glycan trait, and **(E)** G2F glycan trait, were analyzed separately for men, women, and combined groups. The significance of the difference between the slopes of the correlation in PWH and their HIV-negative controls was determined from the interaction term between age and HIV status in a linear regression model.

To validate these cross-sectional findings, we conducted an additional analysis focusing on the longitudinal changes in IgG *N*-glycomic patterns from 23 WWH, 25 MWH, and 47 HIV-negative individuals. For this analysis, we used samples that have been collected from each participant at intervals of one or two years (from age ∼50 to ∼60 years old; **Supplementary Table 3)**, resulting in a total of 6-7 samples per person. All participants were from the cross-sectional population, and they were well-matched in terms of age, race, and BMI at the baseline visit. Using these data, we confirmed several findings from our cross-sectional analysis, despite the shorter age span of the longitudinal analysis. For instance, bisected GlcNAc glycans correlated positively with age in both PWH and HIV-negative controls over time, although the slope of the correlation differed between the two groups; G2F glycan trait correlated negatively with age in both groups; and agalactosylated glycans and the G0FB glycan trait correlated positively with age, but the slope varied between the two groups **(Supplementary Figure 4)**. These findings suggest that HIV and/or ART can alter the pace of age-associated IgG glycomic alterations.

### Machine learning models based on IgG glycans indicate an acceleration of biological aging in PWH

IgG glycans have been used as biomarkers for biological aging in the general population.^18,41^ We, therefore, hypothesized that these IgG glycans could be used to estimate the rate of accelerated biological aging during well-suppressed HIV infection. Prior studies had employed a glycomic clock consisting of multiple individual glycomic features. However, those models were trained on samples from individuals who were not matched for ethnicity and demographic factors to PWH, and thus cannot be directly employed to calculate the biological age of PWH. To address this, we leveraged the analysis of IgG glycomes in HIV-negative individuals from the MWCCS who serve as a comparison group to PWH.^29–32^ Using this data, we developed new glycan-based machine learning models that can estimate the acceleration of biological age in PWH.

We first used the “least absolute shrinkage and selection operator” (LASSO) model to identify the fewest number of individual IgG glycan traits that, when combined, correlated with chronological age in HIV-negative men with an average of zero difference between predicted age and chronological age. This would enable us to calculate the additional rate of acceleration in biological aging in MWH that is specifically influenced by HIV and/or ART. Among the models evaluated (**Table 3**), the best-performing one incorporated four glycan structures: G0, G0FB, G1FB, and A2. When this model was applied to data from MWH, the results (as depicted in **Figure 4A: left**) indicated an average acceleration of 3.52 years with a standard deviation of 7.47 years (*P*<0.0001) between predicted age and chronological age in MWH, compared to their matched HIV-negative counterparts.

**Table 3.** Machine learning models based on IgG glycans and/or inflammatory markers.

| Sex |  | Lasso model |  |  |  | Training model data |  |  |  |  | Testing model |  |  |  |  |  |
| --- | --- | --- | --- | --- | --- | --- | --- | --- | --- | --- | --- | --- | --- | --- | --- | --- |
|  |  | Selected predictor(s) | n | R<sup>2</sup> | Adjusted R<sup>2</sup> | MSE | n | MSE | SD | Low 95% CI | Upper 95% CI | n | MSE | SD | Low 95% CI | Upper 95% CI |
| Men | IgG glycans | G0 + G0FB + G1FB + A2 | 253 | 0.23 | 0.21 | 44.19 | 202 (80%) | 44.22 | 2.22 | 39.88 | 48.57 | 51 (20%) | 45.46 | 9.27 | 27.30 | 63.62 |
|  | IgG glycans | G0 + G0FB + G1FB + A2 | 100 | 0.36 | 0.34 | 38.68 |  |  |  |  |  |  |  |  |  |  |
|  | Inflammatory markers* | CXCL9 + Eotaxin | 100 | 0.02 | 0.00 | 59.42 |  |  |  |  |  |  |  |  |  |  |
|  | IgG glycans + Inflammatory markers | G0 + G0FB + G1FB + A2 + CXCL9 + Eotaxin | 100 | 0.37 | 0.33 | 38.02 |  |  |  |  |  |  |  |  |  |  |
| Women | IgG glycans | A1F + G2F | 235 | 0.31 | 0.31 | 36.25 | 188 (80%) | 36.40 | 1.78 | 32.91 | 39.88 | 47 (20%) | 36.25 | 7.27 | 22.00 | 50.49 |
|  | IgG glycans | A1F + G2F | 100 | 0.25 | 0.23 | 32.16 |  |  |  |  |  |  |  |  |  |  |
|  | Inflammatory markers* | CXCL9 + Eotaxin | 100 | 0.23 | 0.22 | 32.82 |  |  |  |  |  |  |  |  |  |  |
|  | IgG glycans + Inflammatory markers | A1F + G2F + CXCL9 + Eotaxin | 100 | 0.41 | 0.38 | 25.32 |  |  |  |  |  |  |  |  |  |  |

\* Pre-selected MSE = Mean Squared Error SD = Standard Deviation

**Figure 4.**
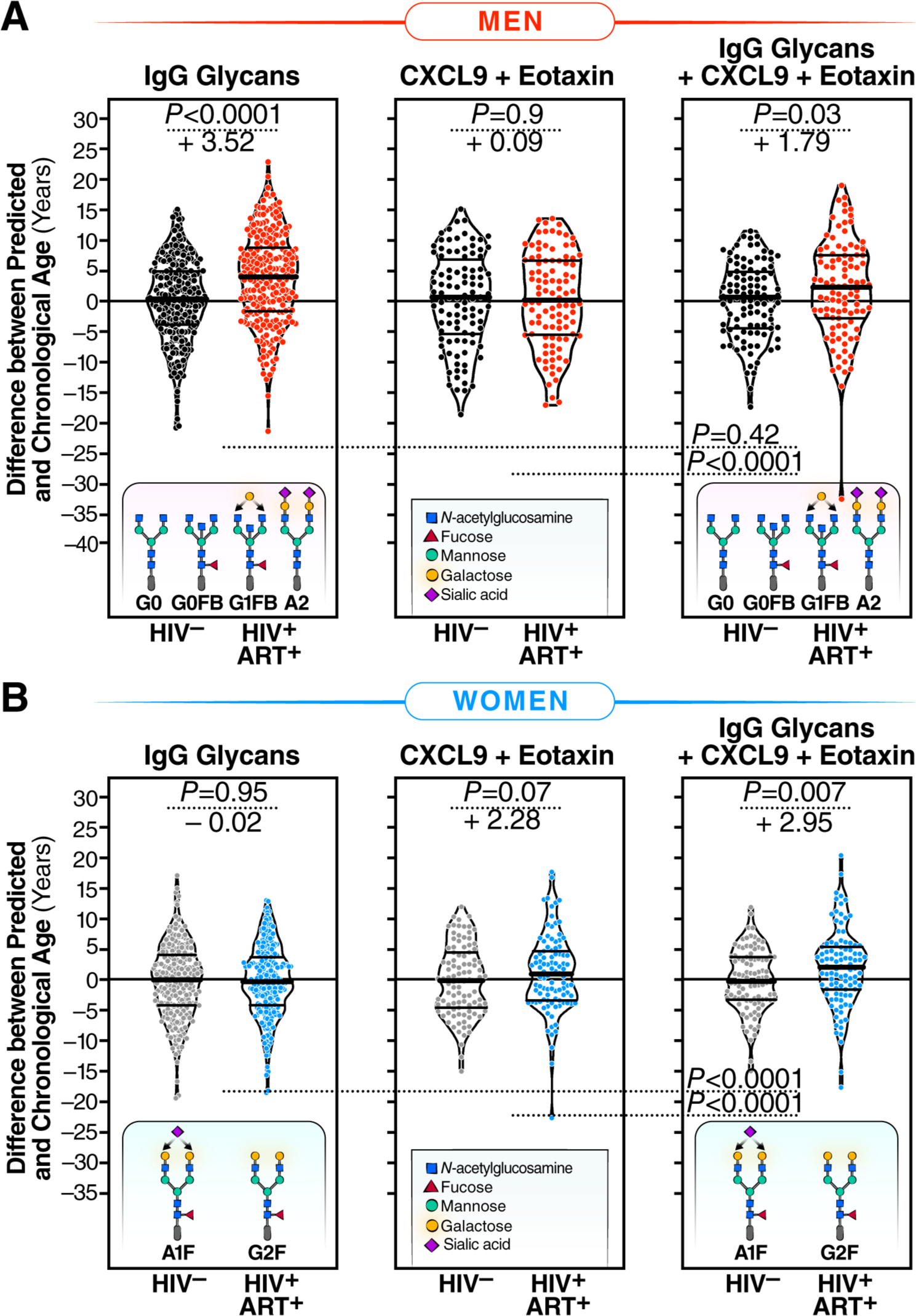
Machine-learning models based on IgG glycans indicate accelerated biological aging during HIV infection suppressed on ART. **(A)** Four specific glycan traits were selected using the LASSO model and included in a model that correlated with chronological age in HIV-negative men, resulting in an average predicted age that closely matched chronological age. In MWH, the model revealed an average acceleration of 3.52 years between predicted and chronological age when compared to their HIV-negative counterparts. The left panel represents the model incorporating only these four glycans. The middle panel includes two inflammatory markers, CXCL9 and Eotaxin. The right panel combines the four glycan traits and the two inflammatory markers. Significance was determined using t-tests. The three models’ efficiency was evaluated using the likelihood ratio test for nested models. **(B)** Two specific glycan traits were selected using the LASSO model and included in a model that correlated with chronological age in HIV-negative women, resulting in an average predicted age that matched chronological age. The left panel represents the model incorporating only these two glycans. The middle panel includes the two inflammatory markers, CXCL9 and Eotaxin. The right panel combines the two glycan traits and the two inflammatory markers. In WWH, only the model combining glycans and inflammatory markers revealed an average acceleration of 2.95 years between predicted and chronological age. Significance was determined using t-tests. The efficiency of the three models was evaluated using the likelihood ratio test for nested models.

As plasma inflammatory markers have also been employed as indicators of biological aging,^40^ we investigated whether models based on some of these markers could also estimate the rate of accelerated biological aging in MWH. We also explored whether the glycomic model’s efficacy could be enhanced by combining plasma inflammatory markers with the four identified IgG glycan structures. First, we examined the correlation between inflammatory markers and chronological age in MWH or WWH (**Supplementary Table 4**). We found that no inflammatory markers correlated with chronological age (FDR<0.05) in MWH, whereas three markers (CXCL9, Eotaxin, and TNFα) did correlate with chronological age (FDR<0.05) in WWH. Next, we assessed whether these three inflammatory markers could be combined in a model. Both CXCL9 and Eotaxin exhibited unique correlations with age; however, their combination was insufficient to estimate the rate of biological aging in MWH (**Figure 4A: middle**; **Table 3**). Furthermore, combining these two inflammatory markers with the four glycans did not improve the predictive capacity of the pure glycan model (**Figure 4A: right**; likelihood ratio test for comparing nested models *P*=0.42; **Table 3**).

Following a similar analysis approach, we identified two glycans (A1F and G2F) in women that correlated with chronological age in HIV-negative women with an average of zero difference between predicted age and chronological age. However, a model based on these two glycans was insufficient in estimating the rate of biological age in WWH (**Figure 4B: left**; **Table 3**). A combination of CXCL9 and Eotaxin moderately estimated the rate of biological aging in WWH (**Figure 4B: middle**; **Table 3**). Notably, the best model for estimating the rate of biological aging in WWH encompassed two glycans (A1F and G2F) and two inflammatory markers (**Figure 4B: right**; *P*<0.0001; **Table 3**). Applying this model to data from WWH, as shown in **Figure 4A: right**, revealed an average acceleration of 2.95 years with a standard deviation of 10.63 years (*P*=0.0066) between predicted and chronological age in WWH.

Taken together, these findings suggest that IgG glycans can be employed in models to calculate biological aging in PWH. However, the model requirements differ between MWH and WWH, whereby glycans alone suffice to estimate the rate of biological aging in MWH, while a combination of glycans and inflammatory aging markers is necessary for similarly robust estimation in WWH.

### IgG agalactosylation and bisecting GlcNAc correlate with the development and severity of coronary atherosclerosis in PWH

We next examined whether the IgG glycomic dysregulations associated with HIV/ART (in Figure 1), that are linked with inflammation (Figure 3), are also associated with the development of age- and inflammation-related comorbidities during ART-suppressed HIV infection. To accomplish this, we first designed a case-control study within the MACS cohort.^42,43^ The cases were individuals with coronary artery stenosis of ≥50% in one or more coronary segments, whereas the controls were individuals with no coronary plaque; as determined by CT angiography. A 1:1 nearest neighbor matching algorithm was employed to select 22 HIV-negative men controls, 22 HIV-negative cases, 34 ART-treated MWH controls, and 34 ART-treated MWH cases (**Figure 5A**). Cases and controls within each group were matched for age, ethnicity, BMI, CD4 T cell counts, and nadir CD4 T cell counts (**Supplementary Table 5**).

**Figure 5.**
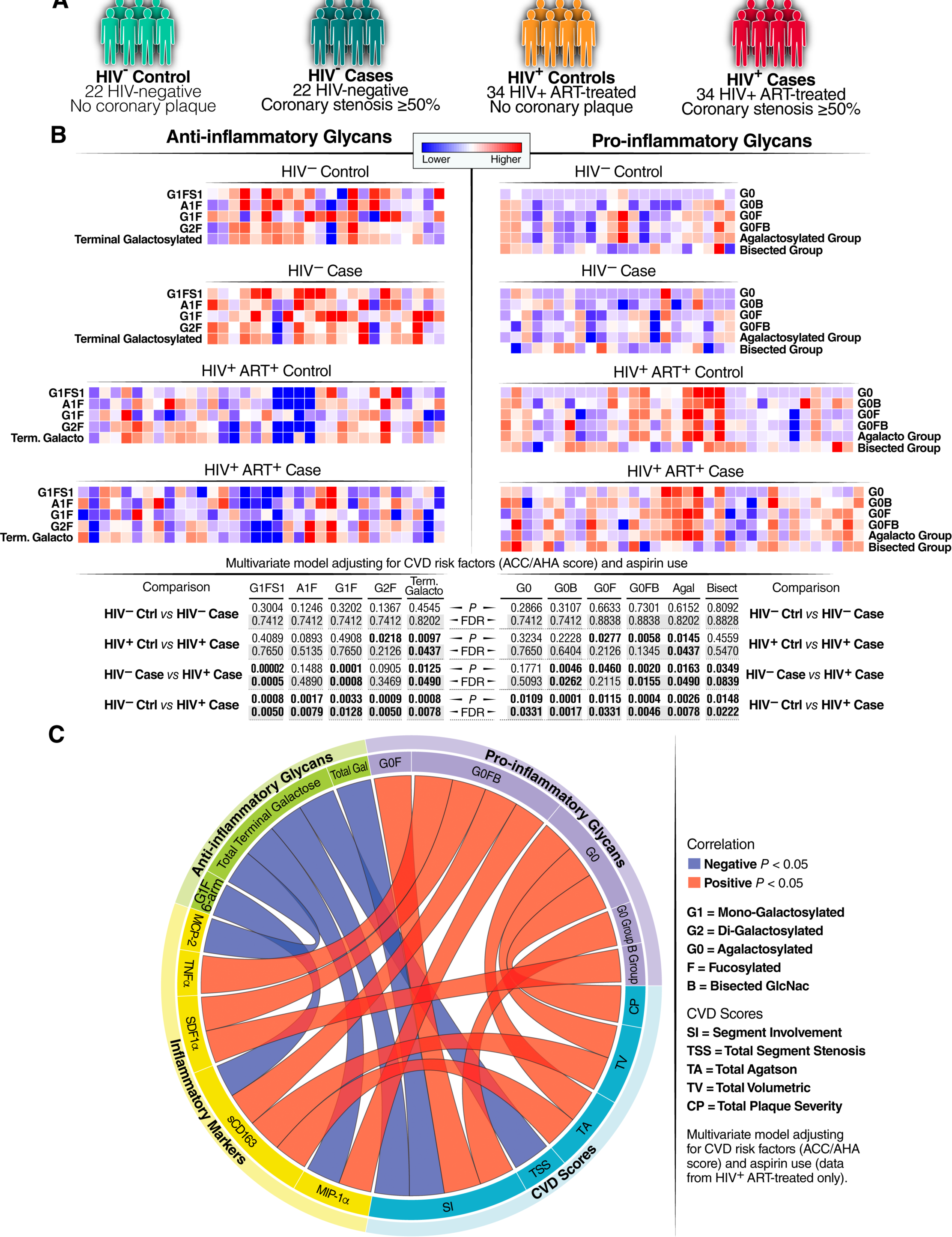
IgG agalactosylation and bisecting GlcNAc correlate with the development and severity of coronary atherosclerosis during ART-treated HIV Infection. **(A)** Schematic representation of the cases and controls included in this analysis. **(B)** Heatmaps illustrating the levels of anti-inflammatory (left) or pro-inflammatory (right) glycans that exhibited statistically significant differences in HIV+ ART+ cases compared to the other three groups. Red indicates higher levels, while blue indicates lower levels. The comparisons were calculated using a linear regression model with independent variables of HIV status, CVD status, and their interaction, adjusting for CVD risk factors (ACC/AHA score) and aspirin use. **(C)** A circos plot displaying Spearman’s rank correlations between anti-inflammatory glycans (green), pro-inflammatory glycans (purple), plasma inflammatory markers (yellow), and CVD scores (teal). Red connections indicate a significant positive correlation, while blue connections indicate a significant negative correlation. Only correlations that remained significant after adjusting for CVD risk factors (ACC/AHA score) and aspirin use are shown. The correlation analysis includes only samples from PWH on ART.

Adjusting for CVD risk factors (ACC/AHA Pooled Cohort Equation risk score) and aspirin use using a multivariable model, we found that levels of specific anti-inflammatory glycan structures, including terminal galactosylated glycans and other galactosylated glycans (such as G2F), were lower in cases living with HIV compared to the other three groups (**Figure 5B: left**). Conversely, levels of several pro-inflammatory glycans, including agalactosylated glycans (the agalactosylated group and individual glycans lacking galactose such as G0, G0B, G0F, G0FB) and the bisected GlcNAc group, were higher in cases living with HIV compared to the other three groups (**Figure 5B: right**; FDR<0.05). Furthermore, levels of many agalactosylated glycans (agalactosylated group, G0, G0F, G0FB) were higher in PWH with coronary plaque compared to those without coronary plaque, while levels of the individual galactosylated glycan trait G2F were lower in PWH individuals with coronary plaque compared to those without coronary plaque (**Supplementary Figure 5**).

We next examined the links between anti-inflammatory or pro-inflammatory glycans, plasma markers of inflammation (**Supplementary Table 6**), and the severity of CVD (measured by various scores depicted in **Figure 5C**). We found that levels of anti-inflammatory IgG glycans, mainly galactosylated glycans, were associated with reduced inflammation and lower CVD severity scores. Conversely, levels of pro-inflammatory glycans, mainly agalactosylated and bisected GlcNAc glycans, were correlated with increased inflammation and higher CVD severity scores (**Figure 5C**). These findings suggest that HIV/ART-mediated alterations in IgG glycomic profiles, particularly the loss of galactose and the gain of bisected GlcNAc, are associated with the development and severity of subclinical coronary artery disease during ART-suppressed HIV infection.

### IgG agalactosylation may precede the development of inflammation-associated comorbidities in PWH

Previous studies in the general population indicated that changes in IgG glycosylation patterns can precede the onset of age-related diseases.^26,27^ Therefore, we conducted an exploratory longitudinal case-control study to investigate whether HIV/ART-related glycomic alterations may precede the development of age-related diseases in PWH on ART. We used longitudinal plasma samples that had been collected from ten PWH on ART before the onset of gastrointestinal (GI) cancers, which are non-AIDS-defining cancers associated with inflammation. Control samples were also obtained from 13 age-, sex-, and ethnicity-matched PWH on ART without cancers at the same time points (**Figure 6A** and **Supplementary Table 7**). Significant differences (FDR<0.05) were observed between cases and controls in the average over time of IgG glycomic groups and individual IgG glycans prior to cancer onset. Specifically, levels of the anti-inflammatory galactosylated and sialylated IgG glycan Groups, as well as several galactosylated individual glycan traits, were lower in cases compared to controls (**Figure 6B, C, D**). Conversely, levels of the pro-inflammatory agalactosylated group and individual glycan traits lacking galactose (such as G0 and G0F) were higher in cases compared to controls, on average, 5-10 years before cancer onset (**Figure 6B, C, E**). Previous research in the general population had identified a specific IgG glycomic ratio called the Gal-ratio, which reflects the distribution of IgG galactosylation and serves as a prognostic biomarker for cancer incidence.^44–46^ In our study, we calculated the Gal-ratio by determining the relative intensities of agalactosylated (G0) versus mono-galactosylated (G1) and di-galactosylated (G2) IgG N-glycans using a previously described formula.^44–46^ Remarkably, we found that the Gal-ratio predicted the occurrence of GI cancer in PWH during ART treatment (**Figure 6B**). This exploratory study suggests that dysregulations in IgG glycomic patterns that are associated with premature aging and with pro-inflammatory responses are not only prevalent during ART-treated HIV infection but also may precede the onset of inflammation-associated diseases in PWH.

**Figure 6.**
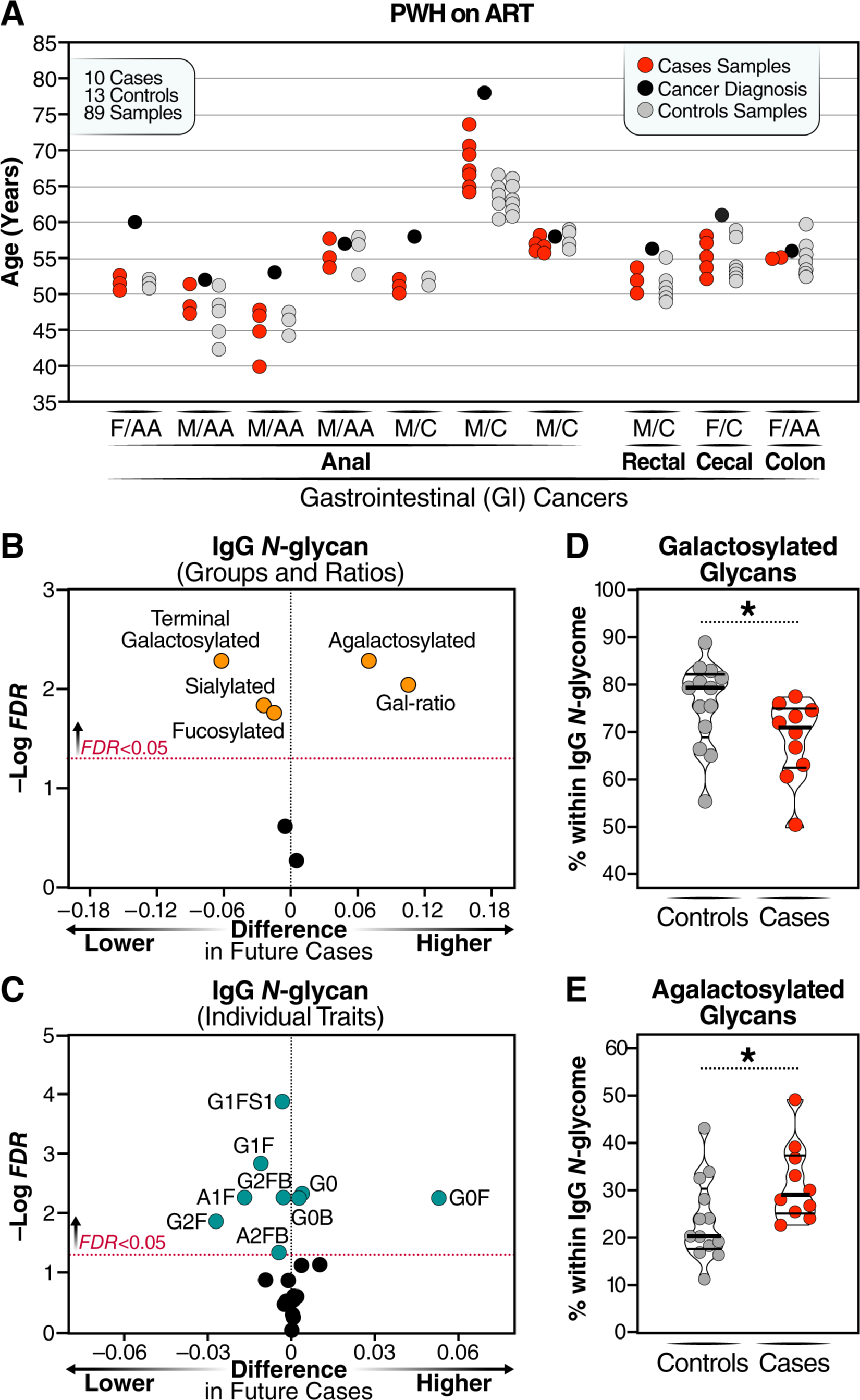
IgG agalactosylation may precede the development of GI-related cancers in PWH on ART. **(A)** Overview of the longitudinal plasma samples analyzed from cases and controls, including the corresponding ages at the time of collection. The labels F and M represent female and male, respectively, while AA and C indicate African American and Caucasian ethnicity. Samples from cases and controls were matched based on age, sex, and ethnicity. **(B-C)** Volcano plots illustrating the IgG glycan groups **(B)** and individual IgG glycans **(C)** that exhibited a significantly higher (right) or lower (left) average over time in cases compared to controls. **(D-E)** Comparison of the average levels of total terminal galactosylated **(D)** and agalactosylated **(E)** IgG over a 5–10 year period before cancer onset in cases, contrasting cases with controls. Median and IQR are displayed. Unpaired t-tests were employed for statistical analysis.

### PWH exhibit heightened expression of senescence-associated glycan-degrading enzymes compared to controls

Our next objective was to investigate the potential upstream mechanisms responsible for the observed HIV/ART-mediated glycomic alterations. Several factors can influence the glycosylation of IgG, including the expression of glycosyltransferases in B cells. Notably, FUT8 catalyzes the transfer of core fucose, MGAT3 catalyzes the transfer of bisected GlcNAc, B4GALT1 catalyzes the transfer of galactose, and ST3GAL1 catalyzes the transfer of sialic acid (**Figure 7A**). On the other hand, the removal of sialic acid and galactose can be attributed to increased systemic levels of glycosidases (NEU1-4 and GLB1, respectively), which can remove these components from IgGs after their production by B cells (**Figure 7A**).^47,48^ We explored these two potential explanations. First, we examined the impact of ART-suppressed HIV infection on the expression of the four aforementioned glycosyltransferases in B cells. To achieve this, we performed single-cell CITE-seq analysis on PBMC from eight PWH on ART and eight HIV-negative controls. Following pre-processing and unbiased clustering to identify immune subpopulations, the B cell cluster was regrouped into five sub-clusters (**Figure 7B**). Upon closer examination, two of these sub-clusters did not express high levels of the B cell markers CD19 and CD20 (**Supplementary Figure 6**) and were therefore excluded from further analysis. We observed modest decreases in the levels of ST3GAL1 and B4GALT1 in B cells from PWH on ART compared to HIV-negative controls, which is consistent with the reduction in levels of IgG sialylation and galactosylation observed in Figure 1 (**Figure 7C**). By contrast, we did not observe any changes in the levels of MGAT3 or FUT8.

**Figure 7:**
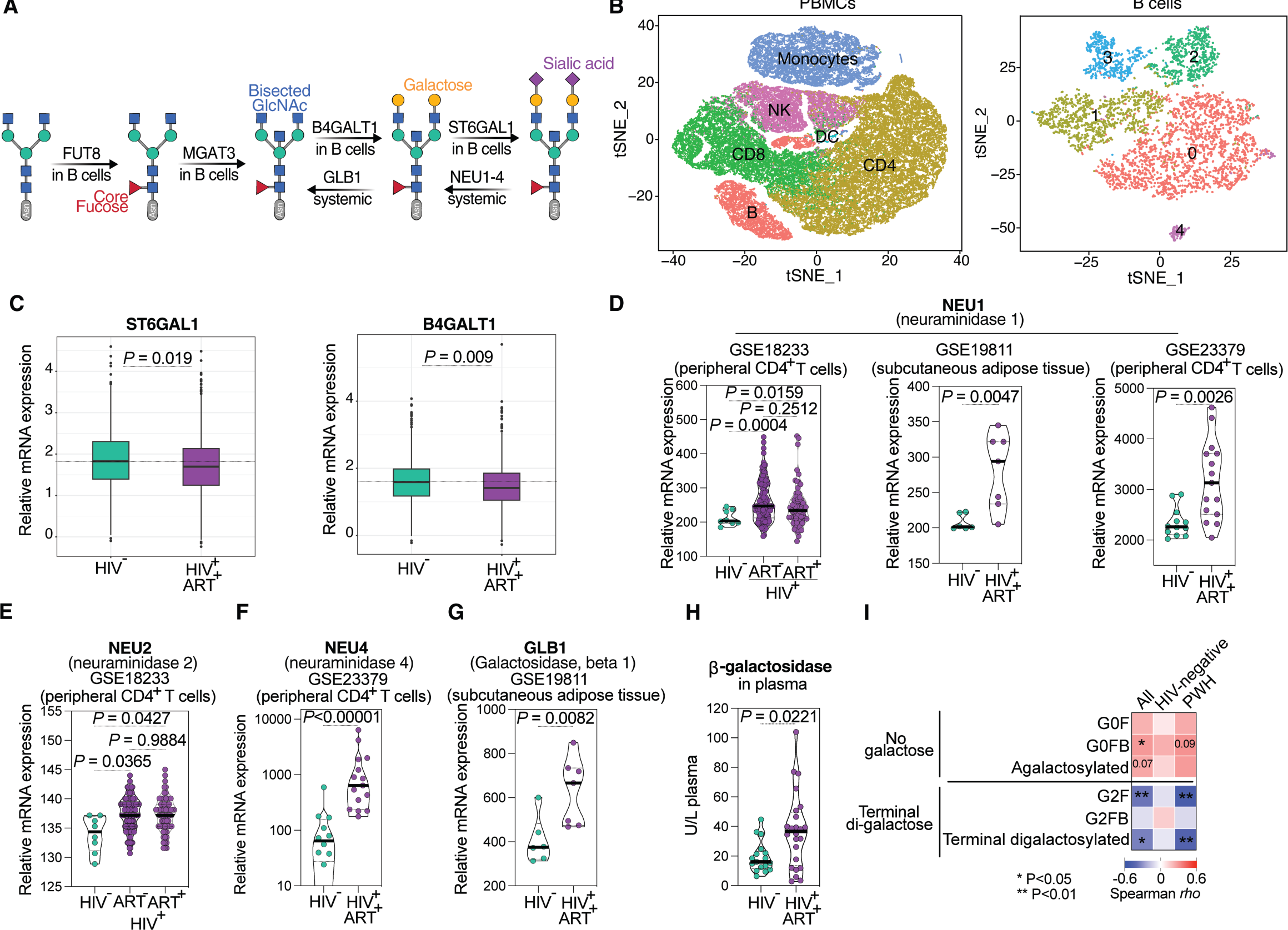
PWH exhibit heightened expression of senescence-associated glycan-degrading enzymes compared to HIV-negative controls. **(A)** Schematic representation illustrating the glycosyltransferases responsible for catalyzing different glycans into IgG glycomes, as well as the glycosidases that remove certain glycans from IgG glycans. **(B)** t-SNE plots displaying PBMCs and B cell clusters from the single-cell cite-seq experiments. **(C)** Single-cell expression levels of ST3GAL1 and B4GALT1 in B cells from PWH on ART and HIV-negative counterparts. Unpaired t-tests were performed for statistical analysis. **(D-G)** Analysis of publicly available gene expression datasets (GSE18233, GSE19811, GSE23379) for genes encoding sialidases (NEU1 (D), NEU2 (E), and NEU4 (F)) and β-galactosidases (GLB1 (G)), comparing their expression levels between PWH and HIV-negative controls. Unpaired t-tests were performed for statistical analysis. ANOVA or Mann-Whitney tests were performed for statistical analysis, and violin plots display the median and IQR. **(H)** Measurement of active β-galactosidase protein levels in the plasma of PWH on ART and HIV-negative controls. Mann-Whitney T-test was performed for statistical analysis, and violin plots display the median and IQR. **(I)** Spearman’s rank correlation heatmap reveals the associations between plasma β-galactosidase in all donors, HIV-negative donors, or PWH on ART, and agalactosylated or galactosylated IgG glycans from the same donors. Positive correlations are depicted in red, while negative correlations are shown in blue. *P<0.05, and **P<0.01.

Next, we explored the second hypothesis that the removal of sialic acid and galactose could also be attributed to increased levels of glycosidases that remove these components from IgGs after their production by B cells. These glycosidases may not necessarily be produced by B cells, but their expression could be systemic. Therefore, we first analyzed gene expression data from various publicly available datasets that examined the transcriptomic profiles of blood and tissue cells from PWH (viremic and suppressed by ART) and HIV-negative controls. These analyses revealed that even after ART suppression, HIV infection was associated with elevated expression levels of several sialidases (enzymes that catalyze sialic acid removal), such as NEU1, NEU2, and NEU4, in peripheral CD4^+^ T cells and cells from adipose tissues compared to controls (**Figure 7D-F**). Additionally, ART-treated HIV infection was linked to increased expression levels of genes encoding β-galactosidase (GLB1, an enzyme that catalyzes galactose removal) in tissues compared to controls (**Figure 7G**). The observation of elevated β-galactosidase mRNA levels in individuals receiving ART for HIV infection was intriguing, as β-galactosidase (β-Gal) is a well-established marker of senescence and aging.^49–51^ Therefore, we sought further to validate the elevation of β-galactosidase during ART-treated HIV infection, and we measured β-galactosidase protein activity in the plasma of 17 HIV-negative individuals and 24 PWH on ART. The results shown in **Figure 7H** indeed demonstrated elevated active β-galactosidase protein levels during ART-treated HIV infection compared to controls. Consistently, levels of active β-galactosidase protein correlated with higher levels of agalactosylated IgG glycans and lower levels of di-galactosylated IgG glycans, especially in PWH (**Figure 7I**). Together, these findings suggest that decreased levels of certain glycosyltransferases in B cells and increased levels of systemic glycosidases, namely sialidase and β-galactosidase, may contribute, at least partially, to some of the pro-inflammatory HIV/ART-associated glycomic alterations.

### HIV/ART-associated IgG glycomic alterations compromise HIV-specific, IgG Fc-mediated antiviral innate immune functions

Our final objective was to explore some of the potential downstream consequences of HIV/ART-induced IgG glycomic alterations, which may mechanistically explain their association with worse clinical outcomes. We specifically aimed to assess whether these glycomic alterations can influence IgG-mediated antiviral activity. In addition to viral antigen neutralization, IgG, through its Fc domain, can engage various innate immune cells to trigger anti-viral innate immune functions. For instance, IgG can engage natural killer (NK) cells to induce ADCC, phagocytic cells to elicit ADCP, and complement proteins to elicit ADCD (**Figure 8A**). The ability of an antibody to trigger these functions depends on several factors, including Fc glycosylation.^10,52–54^ Extensive evidence has shown that the sugar fucose significantly reduces the ability of IgG to induce ADCC.^55^ However, the role of galactose, or the lack thereof, in anti-HIV-specific innate immune functions remains unknown.

**Figure 8.**
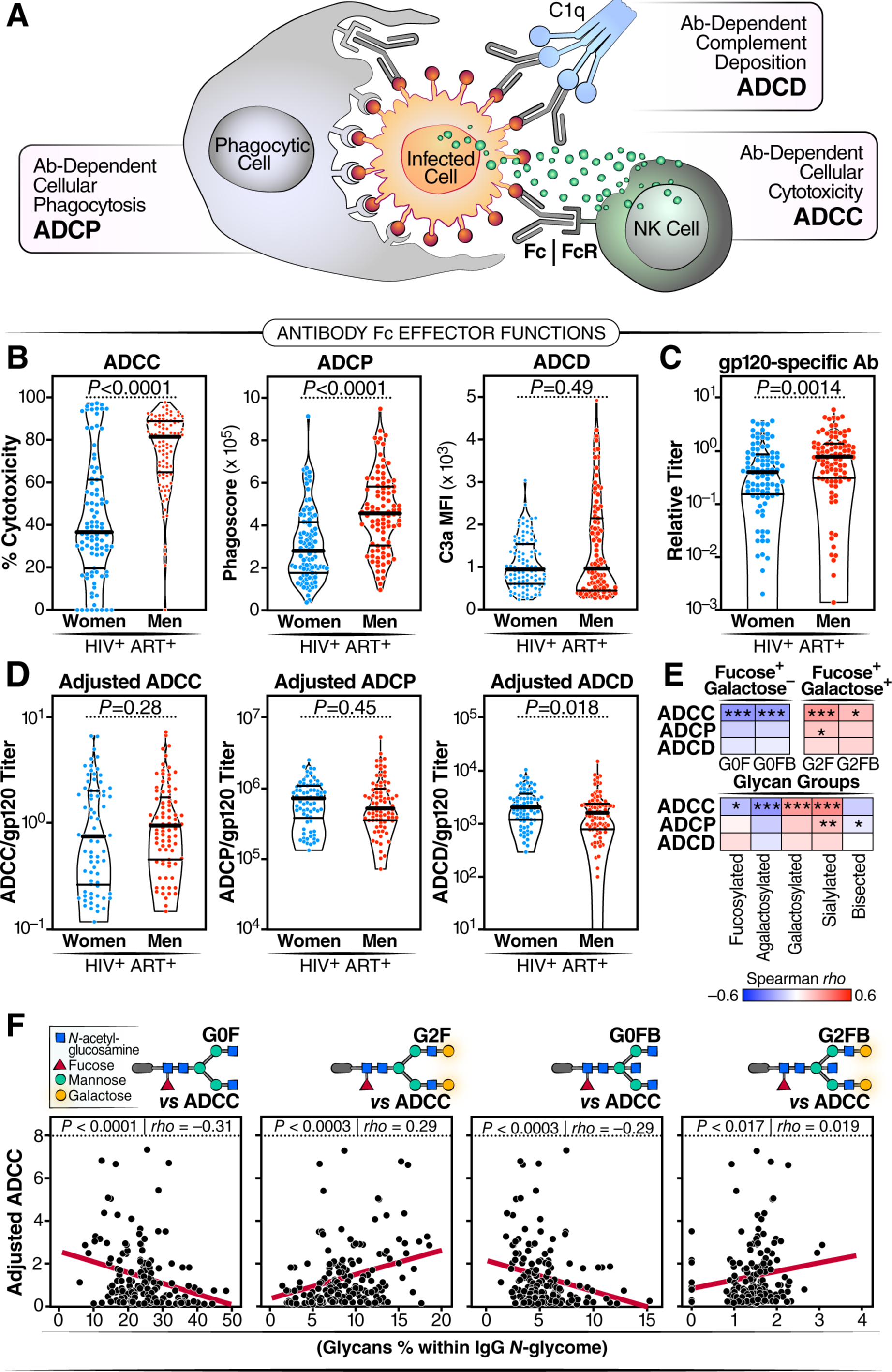
HIV/ART-mediated IgG glycomic alterations are linked to compromised HIV-specific IgG Fc-mediated antiviral innate immune functions. **(A)** Schematic illustration highlighting the role of IgG Fc in facilitating various anti-viral innate immune functions, including ADCC, ADCP, and ADCD. **(B)** Evaluation of ADCC, ADCP, and ADCD elicited by 20µg of bulk IgG from WWH and MWH after background subtraction using samples from HIV-negative men and women, respectively. **(C)** Measurement of gp120-specific antibody levels in the same amounts of bulk IgG from WWH and MWH. **(D)** Normalized data of ADCC, ADCP, and ADCD relative to the levels of gp120-specific antibodies measured in the same amounts of bulk IgG from WWH and MWH. Violin plots display median and IQR, and statistical significance was determined using unpaired t-tests. **(E)** Spearman’s rank correlation heatmaps demonstrate the associations between IgG glycans (columns) and adjusted ADCC, ADCP, and ADCD (rows). Positive correlations are represented in red, while negative correlations are depicted in blue. * P<0.05, ** P<0.01, and *** P<0.001. **(F)** Examples from Spearman’s rank correlations in (E) illustrate that fucosylated glycans lacking galactose exhibit negative correlations with adjusted ADCC, while fucosylated glycans containing galactose display positive correlations with adjusted ADCC.

To investigate this, we selected samples from MWH and WWH and examined their ability to elicit anti-HIV specific ADCC, ADCP, and ADCD using *in vitro* functional assays. Initially, we compared the levels of ADCC and ADCP induced by equal amounts of bulk IgG (20µg) from MWH and WWH, finding that bulk IgG from MWH exhibited higher levels of ADCC and ADCP than did bulk IgG from WWH (**Figure 8B**). However, since these results could be influenced by the quantity of HIV-specific antibodies in this amount of bulk IgG, we measured levels of gp120-specific antibodies in these 20µg samples and found that MWH had higher levels of gp120-specific antibodies than WWH (**Figure 8C**). Therefore, we normalized the ADCC, ADCP, and ADCD abilities of bulk IgG based on the quantity of HIV-specific antibodies within these samples and found no differences between MWH and WWH in ADCC and ADCP. However, ADCD activity was lower in MWH compared to WWH (**Figure 8D**).

Next, we examined the correlations between IgG glycosylation and the normalized ability of IgG to induce ADCC, ADCP, and ADCD. Considering that fucose plays a significant role in IgG-mediated innate immune functions, we focused on glycan structures that contained fucose, with and without galactose. As shown in **Figure 8E-F**, fucosylated agalactosylated glycans exhibited negative correlations with ADCC. However, when the same fucosylated glycans were galactosylated, the correlations became positive, suggesting that galactosylation may improve these innate immune functions, and loss of galactose may compromise these functions. Indeed, total agalactosylated glycans (induced by HIV/ART) negatively correlated with ADCC, while the galactosylated glycans (depleted by HIV/ART) positively correlated with ADCC. Additionally, the sialylated glycans (which are depleted by HIV/ART) correlated positively with ADCC and ADCP, and the bisected GlcNAc glycans (which are induced by HIV/ART) correlated negatively with ADCP (**Figure 8E**).

Beyond the correlations between antibody glycosylation and Fc-mediated innate immune functions, and to unequivocally demonstrate a mechanistic link between the loss of galactose and lower anti-HIV Fc-mediated innate immune functions, especially ADCC as shown in Fig. 8E-F, we used a chemoenzymatic method to engineer IgG glycans. We applied this method to the HIV broadly neutralizing antibody (bNAb) 10-1074, generating four distinct glycoforms (**Fig. 9A**-**B**). Since fucose is known to dramatically reduce ADCC,^55^ we designed these glycoforms so that two of them retain fucose, but one contains galactose (G2F), while the other lacks galactose (G0F). Meanwhile, the other two glycoforms lack fucose, but one contains galactose (G2), while the other lacks galactose (G0). This strategy allowed us to examine if galactose can alter ADCC, beyond the impact of fucose.

**Figure 9.**
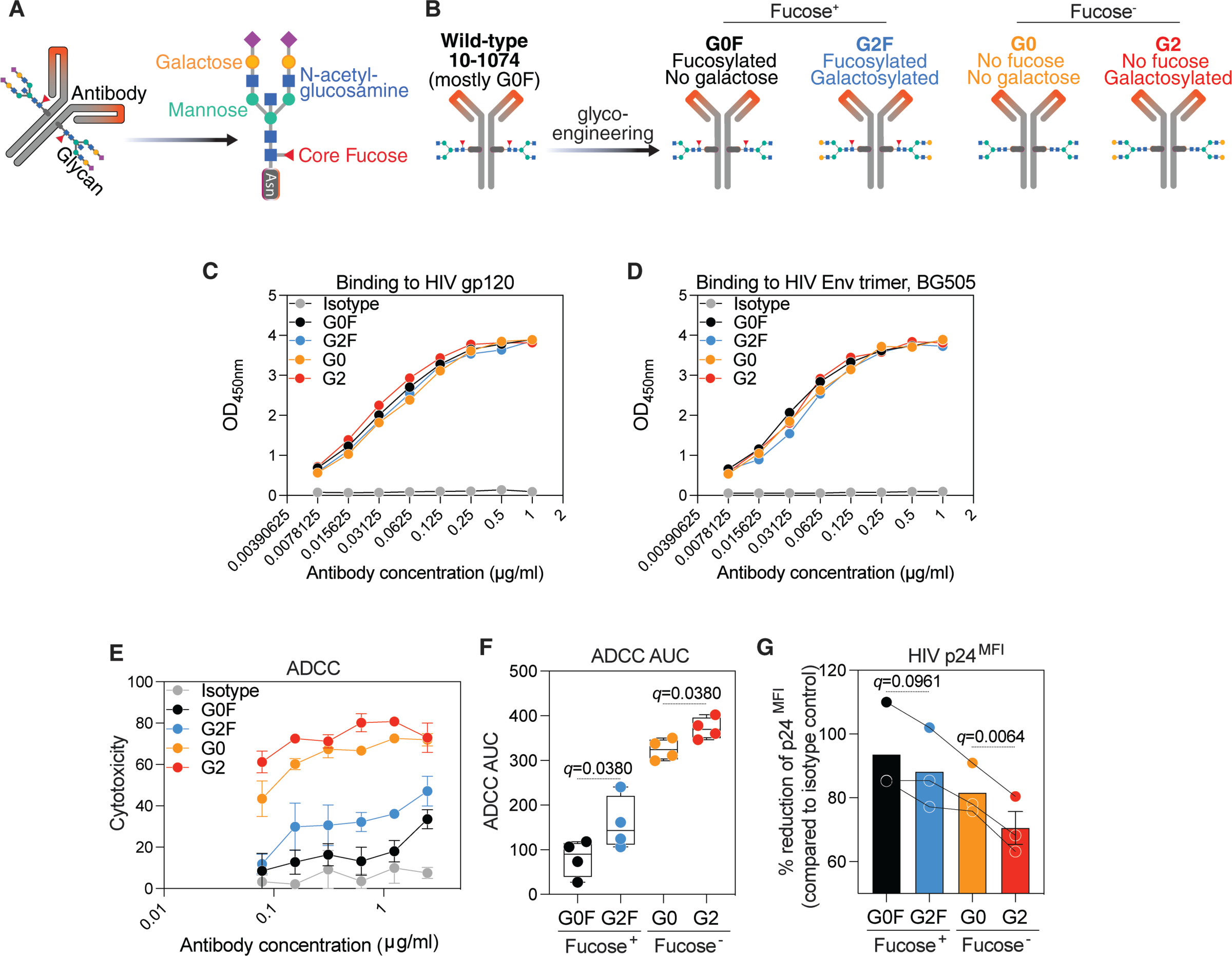
Glycoengineering reveals that IgG agalactosylation reduces anti-HIV ADCC, while galactosylation enhances it. **(A)** Schematic representation of IgG glycans. **(B)** Schematic illustration: 10-1074 was engineered by introducing glycans with four distinct glycomic structures. **(C-D)** The binding of the four glycoforms to HIV-1 gp120 (C) and HIV-1 Env trimer, BG505 (D), assessed by ELISA. Mean and standard error of the mean (SEM) are displayed. **(E-G)** Among fucosylated glycoforms, the galactosylated glycoform (G2F) exhibited higher ADCC **(E-F)**, resulting in lower levels of HIV p24 in infected cells **(G)** compared to the agalactosylated glycoform (G0F). Among afucosylated glycoforms, the galactosylated glycoform (G2) exhibited higher ADCC **(E-F)**, resulting in lower levels of HIV p24 in infected cells **(G)** compared to the agalactosylated glycoform (G0). Analysis in panels E-F was done using one-way ANOVA corrected by the two-stage Benjamini, Krieger, & Yekutieli method. AUC = Area Under the Curve.

We first examined the purity of the four glycoforms using both capillary electrophoresis and mass spectrometry (glycomics and glycoproteomics). As shown in **Supplementary Figure 7**, all four glycoforms are of high purity (>90%). In addition to their purity, these four glycoforms exhibit comparable binding to HIV antigens (**Fig. 9C**-**D**). However, they induce different innate functions: among the fucosylated glycoforms, the glycoform containing galactose (G2F) demonstrated stronger ADCC than the corresponding glycoform lacking galactose (G0F). Similarly, among the non-fucosylated glycoforms, the glycoform containing galactose (G2) demonstrated stronger ADCC than the corresponding glycoform lacking galactose (G0) (**Fig. 9E**-**F**). Consistently, the galactosylated glycoforms enable more efficient ADCC-mediated reduction of the levels of HIV p24 in the cultures compared to non-galactosylated glycoforms, within both fucosylated and non-fucosylated glycoforms (**Fig. 9G**). It is worth noting that the glycosylation and function of wild type 10-1074, currently used in HIV cure clinical trials, are similar to the weak G0F glycoform (**Supplementary Figure 7**). These data suggest that the loss of galactose reduces anti-HIV Fc-effector functions, which is consistent with our *in vivo* data showing that this loss of galactose is associated with worse clinical outcomes and higher inflammation in PWH.

## DISCUSSION

This study is the first comprehensive and controlled investigation examining the relationships between bulk IgG glycomes, chronic inflammation, biological aging, and the development of comorbidities in PWH effectively managed by ART. Our findings suggest that chronic HIV infection, even when suppressed with ART, disrupts the normal progression of age-associated IgG glycomic alterations. These disruptions are linked to increased inflammation, accelerated biological aging, and heightened susceptibility to age- and inflammation-associated comorbidities. These findings have the potential to pave the way to identify novel glycomic-based biomarkers of inflammaging during HIV infection. Furthermore, they can offer valuable insights into the underlying mechanisms driving age- and inflammation-associated diseases in PWH.

While HIV/ART mediate significant alterations in IgG glycans, we have observed a wide range of IgG glycomic profiles among PWH and their controls. IgG glycomic profiles can be influenced by various demographic, environmental, and clinical factors, including genetics, diet, age, sex, gender, pregnancy, smoking status, chronic conditions (such as diabetes), and medications^11,18,33,56–58^ For instance, we observed a significant influence of menopause status on the HIV/ART-mediated IgG glycomic alterations. Future investigations should examine the impact of each of the aforementioned factors, as well as CMV infection status, ART regimen, ART duration, and gender, on IgG glycomes during ART-suppressed HIV infection. This may help identify specific clusters of individuals with distinct glycomic profiles. In our exploratory analysis, we also found that IgG glycomic alterations may precede the development of non-AIDS-defining cancers in PWH on ART. However, this observation requires validation in larger studies with longitudinal samples from PWH who have documented comorbidity outcomes. Such validation, along with the aforementioned clustering analysis, could establish the foundation for using these IgG glycomic profiles individually or in combination (as in a model) as biomarkers to predict the development of aging- and inflammation-associated comorbidities. Such biomarkers may contribute significantly to improving the clinical management of individuals living with chronic viral infections.

In addition to serving as biomarkers of accelerated biological aging and of development of aging-and inflammation-associated diseases,^18,59–63^ IgG glycans are biologically-active molecules that can play significant roles in mediating immunological functions. For instance, they have the ability to modulate the Fc-mediated innate immune functions of antibodies, including ADCC and several pro- and anti-inflammatory activities.^13,64,65^ During ART-suppressed HIV infection, one of the most notable alterations in IgG glycosylation is the loss of galactose (agalactosylation), which correlated with increased inflammation, incidence and severity of subclinical atherosclerosis, and even the development of inflammation-associated cancers in PWH on ART. The causal links between IgG galactosylation and inflammation are well-established and have been proposed to be associated with the ability of IgG galactose to facilitate the interaction between FcγRIIB (CD32b) and dectin-1, leading to anti-inflammatory cascades.^39,66,67^ Conversely, agalactosylation of IgGs (as observed during ART-suppressed HIV infection) has been linked to pro-inflammatory functions by preventing these anti-inflammatory cascades.^39,66,67^

Beyond its inflammatory effects, IgG galactosylation may enhance the activities of antibodies involved in ADCC, ADCP, and CDC.^68,69^ Our data support the possibility that the loss of galactose is associated with and mechanistically can lead to reduced anti-HIV IgG Fc-effector functions, which in turn may contribute to inadequate control of virally-infected cells expressing antigens in tissues. The effectiveness of ART in completely suppressing viral replication, particularly in tissues where ART penetration may be sub-optimal, remains unclear.^70^ It can be hypothesized that the compromised anti-HIV innate immune functions resulting from agalactosylation could contribute to higher HIV persistence and consequently greater inflammation. Further supporting this hypothesis are our previously observed negative correlations between the degree of bulk IgG galactosylation and the levels of cell-associated HIV DNA and RNA in CD4^+^ T cells during ART-suppressed HIV infection.^71^ Further experiments will be needed to determine whether the loss of IgG galactosylation, both in bulk antibodies and antigen-specific antibodies, is directly linked to higher inflammation through the effects of agalactosylation on the complement system or is indirectly linked through the decreased antiviral activity of the antibodies. These studies could help guide the optimization of glycosylation designs of broadly neutralizing antibodies that are currently being explored for HIV cure and prevention strategies. Glycoengineering techniques, such as gene editing and metabolic inhibitors of glycosylation, have been employed to modulate antibody interactions with Fc receptors and enhance their ADCC activity.^72–80^ Understanding the specific glycomic traits that impact HIV or immune functions may pave the way for innovative glycan-based strategies to enhance immune functions during ART-suppressed HIV infection.

There are two potential explanations for IgG glycomic alterations mediated by HIV and/or ART. The first explanation is based on research suggesting that HIV infection leads to certain irreversible defects in B cells, even with ART. These defects may involve changes in the transcriptional profile of glycosyltransferases,^81^ which can result in differential antibody glycosylation. Notably, our previous studies showed that type-I interferons can induce elevated levels of bisected GlcNAc on bulk IgG during ART-suppressed HIV infection.^82^ Cytokines, such as interferons, which are known to play a significant role in glycomic alterations associated with inflammatory diseases,^83^ could also be involved in modulating the glycosylation machinery of tissue plasma B cells, antibody glycosylation, and consequently, the functions of antibodies during ART-suppressed HIV infection. Further investigation is warranted to better understand the potential role of the cytokine milieu in this process during ART-suppressed HIV infection.

The second explanation for the changes in IgG glycomic patterns observed in HIV/ART-mediated conditions is based on research suggesting that there is an increased presence of glycosidases that target sialic acid and galactose, and that these remove glycans from IgGs after the antibody is produced by B cells.^47,48^ Our data support this hypothesis, as we observed higher expression levels of sialidase and β-galactosidase during ART-suppressed HIV infection. This observation that β-galactosidase mRNA levels are elevated in individuals receiving ART for HIV infection is intriguing for two reasons. Firstly, this increase in glycosidases may provide an explanation for the agalactosylation associated with HIV that we observed in our study. Secondly, β-galactosidase is a well-established marker of senescence and aging,^49–51,84–86^ which further reinforces the connection between HIV infection, accelerated aging, and the modulation of glycomic patterns. It is possible that increased levels of glycosidases, which lead to reduced levels of the anti-inflammatory glycans sialic acid and galactose, could be a host response to the virus. That is to say, that the host may upregulate sialidase and β-galactosidases as a mechanism to decrease the anti-inflammatory effects of sialic acid and galactose, thereby promoting inflammation and enhancing the immune response to the infection. Additionally, this response of inducing glycan-degrading enzymes could be triggered by the chronic exposure to microbes from the gut microbiome, which occurs during HIV infection and is associated with translocation of microbes from the gut to the bloodstream. It has been demonstrated that myeloid cells express high levels of endogenous sialidases upon stimulation of Toll-like receptor 4 (TLR4) by bacterial lipopolysaccharide (LPS).^87^ These sialidases remove sialic acids from the surface of the myeloid cells, thereby reducing the potential for inhibitory interactions between Siglec receptors and sialic acids, which could dampen myeloid cell function.^87,88^ While these mechanisms are likely important for maintaining a functional immune response to pathogens, they may also contribute to a state of inflammation. In support of this hypothesis, the loss of galactose from IgG glycomes is not unique to HIV infection, and has been observed during other viral infections and conditions that involve microbial translocation, such as severe SARS-CoV-2 infection.^89–95^

Our study has limitations: 1) Since our study used samples from individuals with ART-suppressed HIV infection, we cannot examine the direct impact of HIV itself in the absence of ART as well as other differences between groups that could serve as confounding variables. However, our previous studies demonstrated similar alterations in IgG glycans in HIV viremic individuals,^71^ suggesting that, at least, some of the alterations observed in this study are mediated by the infection itself. Nonetheless, further research is required to determine if ART toxicity contributes to glycomic alterations, and this can be investigated, at least in part, using samples from Pre-Exposure Prophylaxis (PrEP) trials. Additionally, future studies should investigate whether early initiation of ART can prevent irreversible HIV-associated glycomic alterations. 2) Our focus in this study was on women and men aged ∼40 to over 65 years old. It is important to explore the impact of ART-suppressed HIV infection on younger and older individuals, particularly considering the potential influence of sex hormones in modulating IgG glycans. 3) It will be also necessary to investigate the effect of ART-treated HIV infection on glycomic patterns in diverse populations, e.g., from various geographical regions, including low- and middle-income countries, as well as from areas with different HIV subtypes. 4) Our exploratory analysis in Figure 6 suggests that IgG glycomic alterations may precede the development of inflammation-associated diseases. However, it is essential to validate this possibility by conducting adequately powered longitudinal studies. 5) Detailed investigations into the causes and consequences of these IgG glycomic alterations are necessary. These studies should encompass the analysis of the impact of HIV infection, ART, sex hormones, and inflammatory cytokines on the glycosylation machinery of tissue B cells and the activity of glycosidases in different immune cells. Moreover, it is crucial to examine the influence of glycomic modulation on the Fc-mediated innate immune functions and inflammatory activities of antibodies *in vivo*, which can be accomplished through the use of animal models of HIV infection. Despite these limitations, our current study represents a critical initial step towards elucidating the glycomic mechanistic underpinnings of HIV-associated chronic inflammation in aging population.

## METHODS

### Ethics and study cohorts

The study protocols were approved by the Institutional Review Board of The Wistar Institute, the University of Pennsylvania (IRB protocol 21808309), and the institutions representing the individual MWCCS clinical sites (https://statepi.jhsph.edu/mwccs/). All human experimentation was conducted per the guidelines set forth by the US Department of Health and Human Services and the authors’ respective institutions. Detailed demographic and clinical information regarding the different study cohorts can be found in the results section, as well as in **Table 1** and **Supplementary Tables 3, 5, and 7**. Menopause status was self-reported and classified based on the Stages of Reproductive Aging Workshop 10 years after the first workshop (STRAW+10) criteria: 1) Premenopause (regular menstruation or menstruation with no persistent change that would meet subsequent criteria for peri- or postmenopause); 2) Perimenopause, including early perimenopause (defined as a persistent difference of seven days or more in the length of consecutive cycles) and late perimenopause (occurrence of amenorrhea of 60 days or longer but less than 12 months); and 3) Postmenopause (following 12 months of amenorrhea).

### IgG N-glycan analysis

IgG was purified from 50 µl of plasma using the Pierce Protein G Spin Plate (Thermo Fisher catalog #45204). *N*-glycans were released using peptide-N-glycosidase F (PNGase F) and labeled with 8-aminopyrene-1,3,6-trisulfonic acid (APTS) using the GlycanAssure APTS Kit (Thermo Fisher, catalog #A33952), following the manufacturer’s protocol. The labeled *N*-glycans were analyzed using the 3500 Genetic Analyzer capillary electrophoresis system. The relative abundance of *N*-glycan structures was quantified by calculating the area under the curve of each glycan structure divided by the total glycans using the Applied Biosystems GlycanAssure Data Analysis Software Version 2.0.

### Measurement of plasma inflammatory markers

Plasma levels of Fractalkine, IFN-α2a, IL-12p70, IL-2, IL-4, IL-5, IP-10, MCP-2, MIP-1α, SDF-1α, Eotaxin, IFN-β, IFN-γ, IL-10, IL-1β, IL-21, IL-6, Leptin, CXCL9, and TNF-α were determined using U-PLEX kits from Meso Scale Diagnostics (Biomarker Group 1 (hu) Assays; catalog #K151AEM-2, Custom Immuno-Oncology Grp 1 (hu) Assays; catalog #K15067L-2, and Human MIG Antibody Set; catalog #F210I-3) according to the manufacturer’s instructions. Soluble CD14 and soluble CD163 were measured by ELISA using DuoSet kits from R&D Systems (catalog #DY383 and DY1607, respectively) following the manufacturer’s protocol.

### Measurement of β-galactosidase activity in the plasma

β-galactosidase (GLB1) activity in the plasma was quantified using the Beta-galactosidase Assay Kit (Abbexa; catalog #abx298865) following the manufacturer’s protocol. Briefly, β-galactosidase in the plasma catalyzes the hydrolysis of nitrophenyl-β-galactopyranoside, producing nitrophenol. The concentration of nitrophenol is directly proportional to the enzyme activity in the samples, which can be determined by measuring the absorbance at 400nm.

### Glycoengineering of 10-1074 antibody

The 10-1074 antibody was modified using several kits from Genovis. We utilized the TransGLYCIT™ G0 (Genovis catalog # T1-G0F-010) and TransGLYCIT™ G2 (Genovis catalog # T1-G2F-010) kits to create the glycoforms G2F and G0F from 10-1074. These kits selectively remove only the *N*-glycans on the Fc-part of the IgG, while preserving fucose. After deglycosylation, G0 glycan and G2 glycan were added to the core GlcNAc, followed by IgG purification. To generate glycoforms G2 and G0 without core fucose, we employed the TransGLYCIT™ G2 Afucosylated (Genovis catalog # T1-G2A-010) and TransGLYCIT™ G0 Afucosylated (Genovis catalog # T1-G0A-010) kits. These kits remove both the *N*-glycans on the inner GlcNAc and the α1-6 linked core fucose. G0 glycan and G2 glycan were then added, and IgG was purified in the subsequent step. To assess the purity of these modified glycoforms, we conducted capillary electrophoresis as previously descried. We have also employed two mass spectrometry-based methods: 1) <u>Glycoproteomics</u>: IgGs were reduced with 25 mM dithiothreitol (DTT) at 60°C for 40 minutes. The samples were further alkylated with 90 mM iodoacetamide (IAA) at room temperature for 20 mins. IgGs were desalted by 10 kDa centrifuge cartridges. Trypsin digestion was performed on reduced samples, 1:20/enzyme:protein ratio was used, and material was incubated at 37°C for 18 hours. Prior LC-MS/MS analysis all peptide/glycopeptides were filtered using 0.2 µm filters.^96^ The peptides/glycopeptides were analyzed on an Ultimate 3000 RSLCnano connected to a Thermo Eclipse mass spectrometer. Nano-LC columns of 15 cm length with 75 µm internal diameter, filled with 3 µm C18 reverse phase material were used for chromatographic separation. The separation conditions were low to high acetonitrile in a solution containing 0.1% formic acid, and the separation time was 60 minutes. The precursor ion scan was acquired at 120k resolution in the Orbitrap analyzer and precursors at a time frame of 3 s were selected for subsequent fragmentation using stepped HCD (20, 30 and 40%). Charge state screening was enabled, and precursors with unknown charge state or a charge state of +1 were excluded. Dynamic exclusion was enabled (exclusion duration of 60 s). The centroided fragment ions were analyzed on an Orbitrap detector at 30k resolution. The resulting glycoproteomic data were processed in Byonic v5.2. and pGlyco3 searched against the IgG1 sequence and a custom library of *N*-glycans from human plasma including 24 glycotypes. The precursor mass tolerance and fragment mass tolerances were set to 5ppm and 10ppm respectively. Additional modifications including deamidation of N and Q, carboxymethylation of C, and oxidation of precursor were also included in the search. Assignments were made using Byonic software and manual interpretation. IgG1 glycopeptide assignments was based on the following criteria: Log prob >1, mass error less than 2 ppm (in the most cases), and retention time. Relative abundancies were calculated from the raw data using m/z’s corresponding to the different charge states. 2) <u>Glycomics</u>: 100 µg of the material was reduced with 25 µL of 25 mM DTT and incubated at 50°C for 30 min. After that samples were treated with 90mM iodoacetamide to alkylate. The samples were desalted by centrifugation via Amicon centrifuge filters (10k MWCO size). 3 µL of PNGaseF was added to desalted samples followed by incubation at 37 °C for 18 h to release the N-glycans. The released glycans were permethylated using sodium iodide in the presence of sodium hydroxide and DMSO. Addition of water quenched the permethylation reaction and permethylated *N*-glycans were extracted with dichloromethane. The dichloromethane layer was rinsed five times with water and dried via evaporation by nitrogen gas.^97^ Dried, permethylated glycans were re-dissolved in a solution of 100µL of water and 100µL of methanol for a total volume of 200µL (with 1mM NaOH). Samples were then run on a Themo Orbitrap Fusion Tribrid tandem MS coupled to Ultimate 3000 RSLCnano. 10 μL of *N*-glycans were injected into LC-MS/MS and ran in the low to high organic solvent gradient for 72 minutes. The precursor ion scan was acquired at 120k resolution in the Orbitrap analyzer and precursors at a time frame of 3 s were selected for subsequent fragmentation using CID (45%). Charge states were targeted from 1 to 5. Dynamic exclusion was enabled (exclusion duration of 30s). The centroided fragment ions were analyzed on an Orbitrap detector at 15k resolution. GRITS software and Glyco Workbench 2 was used to identify the *N*-glycan structures (sodium adduct). Relative percentages were retrieved from MS1 level using m/z’s for multiple charge states. FreeStyle 1.8 was used in order to pull out the peaks. Finally, to assess whether the modifications made to the 10-1074 antibody alter its binding to HIV, we performed examined their binding to HIV-1 gp120 and Env trimer BG505 using ELISA.

### Antibody-dependent cellular cytotoxicity (ADCC)

ADCC induction was estimated using CEM.NKR CCR5^+^ Luc^+^ cells co-cultured with primary human NK cells isolated from the blood of healthy donors using negative selection (Stem Cell; catalog #19055) in the presence of purified bulk IgG. HIV-infected and uninfected CEM.NKR CCR5+ Luc+ cells were washed and plated at 2 x 10^4^ cells/well in a V-bottom microplate. The cells were treated with purified bulk IgG from study participants for 2 hours at 37°C. Control wells without antibodies were adjusted to volume with complete medium. After incubation, purified human NK cells were added to the wells, mixed, pelleted at 200g for 2 minutes, and incubated for 16 hours at 37°C. To evaluate luciferase activity, 100 μl of supernatant was removed from all wells and replaced with 100 μl of Bright-Glo luciferase substrate reagent (Promega; catalog #E2610). After 2 minutes, the well contents were mixed and transferred to a clear-bottom black 96-well microplate. Luminescence (RLU) measurements were integrated over one second per well. The raw RLU values are shown relative to the light output generated in the medium-only control (background). For experiments involving glycoengineered glycoforms, following the ADCC assay, cells were collected for intracellular p24 staining. After washing the cells with PBS, they were stained with the following antibodies: anti-CD3 Alexa Fluor 488 (BD Biosciences cat # 557694), anti-CD16 PerCP/Cy5.5 (BioLegend catalog # 302028), anti-CD56 APC/Cy7 (BioLegend catalog # 362512). After staining, cells were fixed with 100 µL of Cytofix (Cytofix/Cytoperm Kit BD catalog # 554714) for 15 minutes at room temperature and permeabilized with 1X Perm/wash buffer. The cells were then incubated for 30 minutes with the following antibodies: anti-HIV-1 core antigen (p24)-RD1 (Beckman Coulter catalog # 6604667), anti-TNFα PE/Dazzle 594 (BioLegend catalog # 502946), anti-IFNγ BV421 (BioLegend catalog # 506538), CD107a PE/Cy7 (BioLegend catalog # 328618). After this incubation, the cells were washed, resuspended, and at least 100,000 events were acquired by flow cytometry using a BD Biosciences FACS Symphony.

### Antibody-dependent cellular phagocytosis (ADCP)

ADCP was measured using a flow cytometry-based phagocytosis assay.^98^ Biotinylated recombinant HIV-1 gp120 from strain CN54 (Acro Biosystems; catalog #GP0-V182E6) was combined with fluorescent NeutrAvidin beads (Life Technologies; catalog #F8776) at a concentration of 1 µg protein per µl bead and incubated overnight at 4°C. The beads were washed twice with 0.1% PBS-BSA to remove excess unconjugated antigens. The gp120-coated beads were resuspended in a final volume of 1 ml in 0.1% PBS-BSA. A 10 µl bead suspension was added to each well of a round-bottom 96-well culture plate, mixed with purified bulk IgG from study participants, and incubated for 2 hours at 37°C. Then, 5 x 10^4^ THP-1 cells in 200 µl growth medium were added to each well and incubated overnight at 37°C. The following day, 100 µl of supernatant from each well was removed, and 100 µl of BD Cytofix™ Fixation Buffer (BD; catalog #554655) was added to each well. Cells were analyzed by flow cytometry, and the data collected were analyzed in FlowJo software. The percentage of fluorescent or bead-positive cells and the median fluorescence intensity of the phagocytic cells were computed to determine a phagoscore.

### Antibody-dependent complement deposition (ADCD)

ADCD was assessed using a flow cytometry-based complement-fixing assay.^99^ Briefly, 10 x 10^6^ HUT78 cells were pulsed with 2 µg of recombinant HIV-1 gp120 from strain CN54 (Acro Biosystems; catalog #GP0-V182E6) for 20 minutes at 37°C. Unbound gp120 was removed by washing the cells twice with 1% PBS-BSA buffer. Bulk IgG samples from study participants were added to the antigen-pulsed cells and incubated for another 30 minutes at 37°C. Freshly resuspended lyophilized guinea pig complement (Cedarlane Labs; catalog #CL4051), diluted 1:20 with veronal buffer 0.1% gelatin with calcium and magnesium (Boston BioProducts; catalog #IBB-300), was added to the cells for 2 hours at 37°C. After washing with 1X PBS, the cells were assessed for complement deposition by staining with goat anti-guinea pig C3-FITC (MP Biomedicals; catalog #0855385). After fixing, the cells were analyzed by flow cytometry, and ADCD was reported as the mean fluorescence intensity (MFI) of FITC-positive cells.

### Measurement of gp120-specific antibodies

gp120-specific antibodies were measured by ELISA. Plates (Nunc maxisorp, flat bottom, Life Technologies; catalog #44240421) were coated with 2.5 μg/mL of gp120 protein from strain CN54 (Acro Biosystems; catalog #GP4-V15227) overnight. The plate was washed with PBS-Tween20 and blocked with SuperBlock™ Blocking Buffer (Thermo Scientific; catalog #37515), and bulk IgG was added. Unbound IgG antibodies were washed, and bound IgG antibodies were detected using HRP-conjugated anti-human IgG antibody. After another wash, the plates were developed with TMB substrate (R&D; catalog #DY999) for 5-10 minutes in the dark, and the reaction was stopped using stop solution (R&D; catalog #DY994). The plates were immediately read at an optical density of 450nm.

### Single-cell RNA-sequencing

Samples were uniquely barcoded using TotalSeq-B human hashtag antibodies (BioLegend, San Diego, CA), as per manufacturer’s directions, to allow for sample multiplexing for the 10x Genomics Chromium Controller single-cell platform (10x Genomics, Pleasanton, CA). Specifically, 2 million PBMC of each sample were first blocked with Human TruStain FcX (BioLegend; catalog #422301), then incubated with 40ng of various anti-human hashtag antibodies carrying unique cell barcodes and 12-100ng of various CITE-seq antibodies, each previously optimized by flow cytometry. TotalSeq-B human CITEseq antibodies (BioLegend) used to subtype immune cells in the downstream data were: anti-human CD14 (#367145), anti-human CD16 (#302063), anti-human CD19 (#302263), anti-human CD20 (#302361), anti-human CD27 (#302851), anti-human CD3 (#300477), anti-human CD38 (#356639), anti-human CD4 (#300565), anti-human CD56 (#362561), anti-human HLA-DR (#307661), anti-human CD45RA (#304161), and anti-human CD8 (#344757). After washing and dead cell exclusion using EasySep™ Dead Cell Removal (Annexin V) Kit (Stem Cell; catalog #17899), a 10x G chip lane was super-loaded with a multiplexed pool of four uniquely barcoded samples, at a total of 40,000 single cells per lane with expected target recovery of 50%, five lanes were loaded in total. Single-cell droplets were generated using the Chromium Next GEM single cell 3’ kit v3.1 (10x Genomics) on the 10x Genomics Chromium Controller. cDNA synthesis and amplification, library preparation, and indexing were done using the 10x Genomics Library Preparation kit (10x Genomics), according to the manufacturer’s instructions. Overall library size was determined using the Agilent Bioanalyzer 2100 and the High Sensitivity DNA assay, and libraries were quantitated using KAPA real-time PCR. Each library consisting of four pooled samples was sequenced on the NextSeq 500 (Illumina) using a 75 base pair cycle sequencing kit (Illumina) and a paired-end run with the following run parameters: 28 base pair x 8 base pair (index) x 55 base pair.

Cell Ranger Suite (v3.1.0, https://support.10xgenomics.com) with refdata-cellranger-GRCh38-3.0.0 transcriptome as a reference was used to map reads on human genome (Hg38) using STAR.^100^ Samples from eight PWH on ART and eight HIV-negative controls were combined and cells expressing fewer than 500 genes and genes that were not expressed in at least one cell were discarded. There were 61,114 cells left after initial quality control. Seurat v3^101^ was used for clustering, marker identification, and visualization. Cell types were predicted against five human datasets as references using SingleR^102^ and a consensus resolution of clusters into six immune cell types was made. Of these, the B cell cluster (5,186 cells) was subset and re-clustered. Gene expression was used to investigate if the sub-clusters were all indeed B cells. Two sub-clusters lacked in expression of known B cell markers and instead expressed genes associated with dendritic cells and T cells, likely due to spill over from neighboring clusters and/or ambient RNA. These were excluded from the analysis. The R package scCustomize was used to visualize gene expression of select markers as estimated kernel densities.

### Statistical analysis

The statistical methods used in data analysis for each figure are described in the corresponding figure legend. In general, two-group t-tests or Mann-Whitney tests were used to determine the difference between any two groups, and false discovery rates (FDRs) were calculated to account for multiple tests over the studied markers. Spearman’s rank correlation analysis was used to evaluate correlations between variables. To determine the difference in the relationship between a studied marker and age, a linear regression model with main effects of age and HIV status, and the interaction term between age and HIV status, was applied. To evaluate the difference in studied markers among the combined conditions of HIV and CVD status, linear regression models were used with independent variables of HIV status, CVD status, and their interaction term, adjusting for CVD risk factors (ACC/AHA score) and aspirin use. To explore which biomarkers of IgG glycomes could be predictors of chronological age using data from HIV-negative individuals, who serve as representative counterparts to PWH in the MACS/WIHS Combined Cohort Study, separately by sex assigned at birth, a linear regression model with the LASSO technique was first carried out using the 5-fold cross-validation (CV) selection option and the one-standard-error rule for determining the optimal tuning parameter. Due to the modest sample size, variable selection was determined using 100 independent rounds of CV LASSO. The biomarkers that were selected 80 or more times out of the 100 runs were used as the final set of predictors in our models.

### Data availability

The authors declare that data supporting the findings of this study are available within the paper and its supplementary information files. Access to individual-level data from the MACS/WIHS Combined Cohort Study Data (MWCCS) may be obtained upon review and approval of a MWCCS concept sheet.LJ Links and instructions for online concept sheet submission are on the study website.

## SUPPLEMENTARY MATERIALS

**Supplementary Figure 1. (A)** Schematic of IgG glycan analysis by capillary electrophoresis. **(B)** Illustration of the various glycan traits included in the IgG glycomic groups.

**Supplementary Figure 2. Impact of menopause/age on IgG glycans during ART-suppressed HIV infection.** Kruskal-Wallis comparisons of IgG glycomic groups between pre-, peri-, and post-menopause women living with or without HIV. Violin plots depict median and IQR.

**Supplementary Figure 3. Impact of menopause/age on inflammation during ART-suppressed HIV infection.** Kruskal-Wallis comparisons of inflammation markers between pre-, peri-, and post-menopause women living with or without HIV. Violin plots illustrate the median and IQR.

**Supplementary Figure 4. Longitudinal analysis of IgG glycans in PWH and HIV-negative controls.** Examples of the analysis of IgG glycans over time in PWH and their age-, sex-, ethnicity-, and BMI-matched HIV-negative controls. The significance of change over time and the difference in change over time between the groups were determined using multivariable mixed effects models.

**Supplementary Figure 5. Association of IgG glycans with the incidence of coronary plaques in PWH on ART.** Unpaired t-test comparisons were performed between individuals living with or without HIV, stratified by the presence or absence of any coronary plaques, to examine the relationship between IgG glycans and plaque incidence.

**Supplementary Figure 6: Expression of key B cell markers in B cell clusters.** t-SNE plots depicting the gene expression densities of CD19, CD20, CD27, and CD38 in the various B cell clusters observed from the single-cell CITE-seq experiments. The two cluster on the top did not express high levels of the B cell markers, CD19 and CD20 and were excluded from the analysis.

**Supplementary Figure 7: Purity of the 10-1074 glycoforms.** Percentage of glycans within each glycoform and wild-type 10-1074 assessed using capillary electrophoresis in triplicate (left), mass spectrometry based glycomics (middle), and mass spectrometry based glycoproteomics (right).

**Supplementary Table 1.** Comparisons of IgG glycans among pre-menopause, peri-menopause, and post-menopause women.

**Supplementary Table 2.** Correlations between IgG glycans and chronological age.

**Supplementary Table 3.** Characteristics of participants in the longitudinal analysis presented in Supplementary Figure 4.

**Supplementary Table 4.** Correlations between inflammatory markers and chronological age.

**Supplementary Table 5.** Baseline characteristics of the subclinical atherosclerosis study participants in studies presented in Figure 5.

**Supplementary Table 6.** Plasma markers of inflammation in the subclinical atherosclerosis study participants in studies presented in Figure 5.

**Supplementary Table 7.** Characteristics of participants in studies presented in Figure 6.

## AUTHOR CONTRIBUTIONS

M.A-M conceived and designed the study. L.B.G carried out the majority of experiments. O.S.A and S.H.L performed the ADCC, ADCP, and ADCD assays. S.S and P.A performed the mass spectrometry based glycomic analyses. T.K and A.K. analyzed the single-cell cite-seq data. Q.L, X.Y, S.L, J.D, D.L, J.Z, and J.L.L.C.A performed statistical analyses. D.B.H, I.O, J.L, M.A.F, S.H, B.M, A.A, B.D.J, C.R, D.M, N.R.R, O.K, S.G, S.W, M.W, W.S.P, A.L, I.F, P.C.T, R.G, and T.T.B selected study participants and interpreted clinical data. L.B.G, Q.L, O.S.A, and M.A-M wrote the manuscript, and all authors edited it.

## ACKNOWLEDGMENTS

This work is mainly supported by the NIH R01AG062383, R01AG062383-04S1, R21AI170166, and the NCI supplement to the Wistar Institute Cancer Center (P30 CA080815) to M.A-M. M.A-M is also funded by the NIH grants, R01AI165079, R01NS117458, R01DK123733, Penn Center for AIDS Research (P30 AI 045008), and the NIH-funded BEAT-HIV Martin Delaney Collaboratory to cure HIV-1 infection (1UM1Al126620). Mass spectrometry based glycomic analyses was partially supported by NIH R24GM137782 and GlycoMIP, a National Science Foundation Materials Innovation Platform funded through Cooperative Agreement DMR-1933525. We Would like to thank Drs. Michel Nussenzweig and Luis J. Montaner for providing the wild-type 10-1074 for the glycoengineering experiments and Dr. Daniel Kulp for providing HIV-1 Env trimer, BG505. We would like to thank Rachel E. Locke, Ph.D., for providing comments.

Data in this manuscript were collected by the MACS/WIHS Combined Cohort Study (MWCCS). The contents of this publication are solely the responsibility of the authors and do not represent the official views of the National Institutes of Health (NIH). MWCCS (Principal Investigators): Atlanta CRS (Ighovwerha Ofotokun, Anandi Sheth, and Gina Wingood), U01-HL146241; Baltimore CRS (Todd Brown and Joseph Margolick), U01-HL146201; Bronx CRS (Kathryn Anastos, David Hanna, and Anjali Sharma), U01-HL146204; Brooklyn CRS (Deborah Gustafson and Tracey Wilson), U01-HL146202; Data Analysis and Coordination Center (Gypsyamber D’Souza, Stephen Gange and Elizabeth Topper), U01-HL146193; Chicago-Cook County CRS (Mardge Cohen and Audrey French), U01-HL146245; Chicago-Northwestern CRS (Steven Wolinsky, Frank Palella, and Valentina Stosor), U01-HL146240; Northern California CRS (Bradley Aouizerat, Jennifer Price, and Phyllis Tien), U01-HL146242; Los Angeles CRS (Roger Detels and Matthew Mimiaga), U01-HL146333; Metropolitan Washington CRS (Seble Kassaye and Daniel Merenstein), U01-HL146205; Miami CRS (Maria Alcaide, Margaret Fischl, and Deborah Jones), U01-HL146203; Pittsburgh CRS (Jeremy Martinson and Charles Rinaldo), U01-HL146208; UAB-MS CRS (Mirjam-Colette Kempf, Jodie Dionne-Odom, Deborah Konkle-Parker, and James B. Brock), U01-HL146192; UNC CRS (Adaora Adimora and Michelle Floris-Moore), U01-HL146194. The MWCCS is funded primarily by the National Heart, Lung, and Blood Institute (NHLBI), with additional co-funding from the Eunice Kennedy Shriver National Institute Of Child Health & Human Development (NICHD), National Institute On Aging (NIA), National Institute Of Dental & Craniofacial Research (NIDCR), National Institute Of Allergy And Infectious Diseases (NIAID), National Institute Of Neurological Disorders And Stroke (NINDS), National Institute Of Mental Health (NIMH), National Institute On Drug Abuse (NIDA), National Institute Of Nursing Research (NINR), National Cancer Institute (NCI), National Institute on Alcohol Abuse and Alcoholism (NIAAA), National Institute on Deafness and Other Communication Disorders (NIDCD), National Institute of Diabetes and Digestive and Kidney Diseases (NIDDK), National Institute on Minority Health and Health Disparities (NIMHD), and in coordination and alignment with the research priorities of the National Institutes of Health, Office of AIDS Research (OAR). MWCCS data collection is also supported by UL1-TR000004 (UCSF CTSA), UL1-TR003098 (JHU ICTR), UL1-TR001881 (UCLA CTSI), P30-AI-050409 (Atlanta CFAR), P30-AI-073961 (Miami CFAR), P30-AI-050410 (UNC CFAR), P30-AI-027767 (UAB CFAR), P30-MH-116867 (Miami CHARM), UL1-TR001409 (DC CTSA), KL2-TR001432 (DC CTSA), and TL1-TR001431 (DC CTSA). The MACS CVD2 study is funded by National Heart Lung and Blood Institute (NHLBI), R01 HL095129-01 (Wendy Post). The authors gratefully acknowledge the contributions of the study participants and dedication of the staff at the MWCCS sites.

