## Supplementary Figure 1 for "Plasma Glycomic Markers of Accelerated Biological Aging During Chronic HIV Infection"

**A**

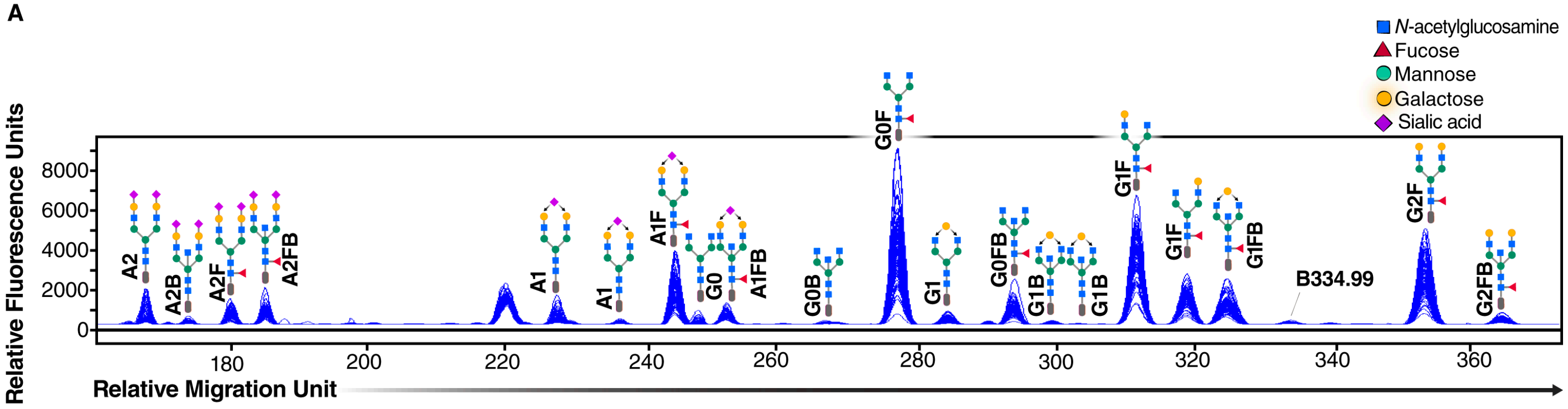


**B**

| IgG Glycosylation group | Glycan traits |
|---|---|
| **Total sialylated group** | A1[3] + A1[6] + A1F + A1FB + A2 + A2B + A2F + A2FB |
| **Agalactosylated group** | G0 + G0B + G0F + G0FB |
| **Total terminal galactosylated** | A1[3] + A1[6] + A1F[3] + A1F[6] + G1 + G1B[3] + G1B[6] + G1F[6] + G1F[3] + G1FB + G2F + G2FB |
| **Fucosylated group** | A2F + A2FB + A1F + A1FB + G0F + G0FB + G1F[6] + G1F[3] |
| **Bisected Group** | A2B + A2FB + A1FB + G0B + G0FB + G1B[3] + G1B[6] + G1FB + G2FB |
