## Supplementary figures and images for "Plasma Glycomic Markers of Accelerated Biological Aging During Chronic HIV Infection"

### Supplementary Figure 2

#### Supplementary Figure 2

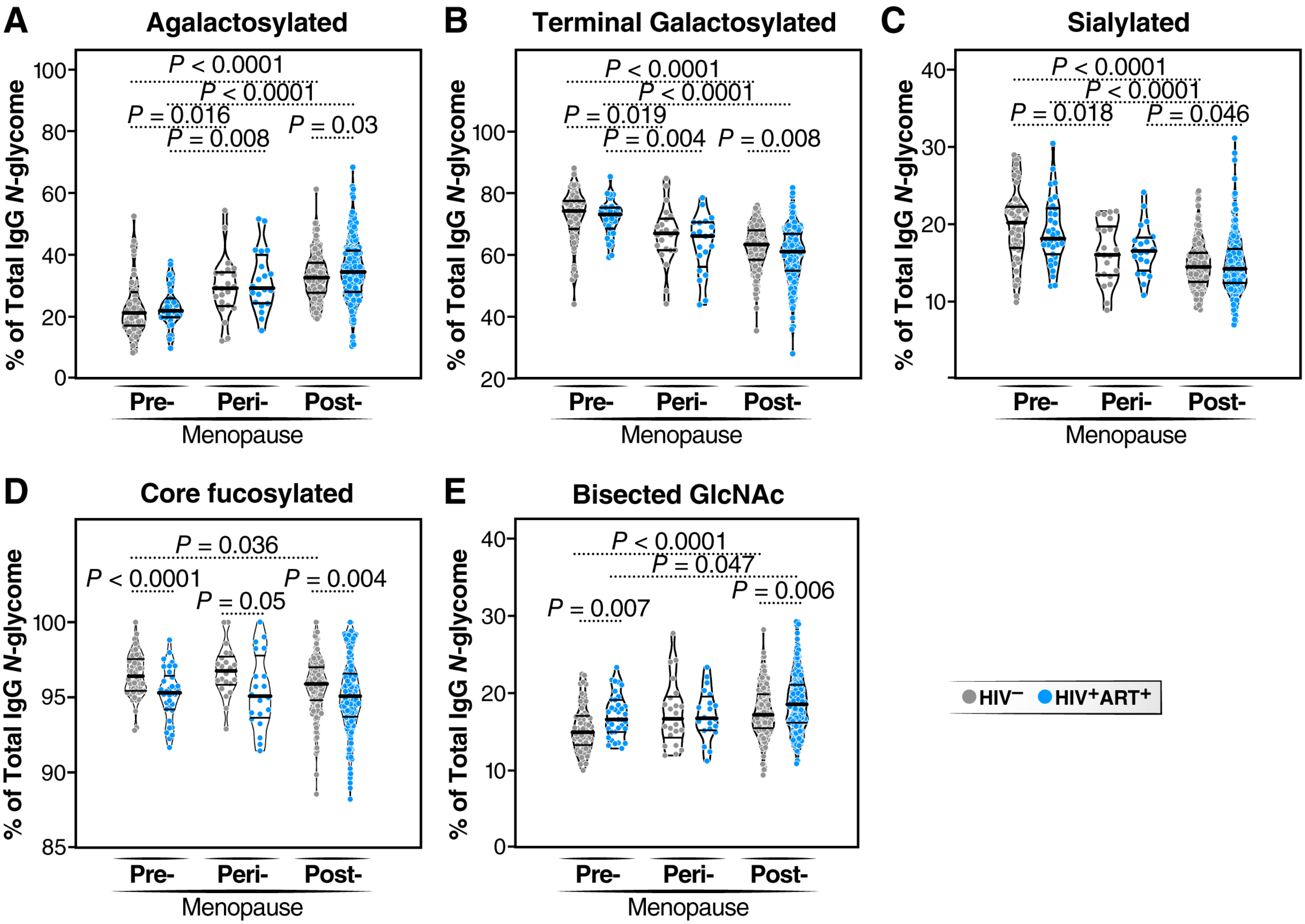

### Supplementary Figure 3

#### Supplementary Figure 3

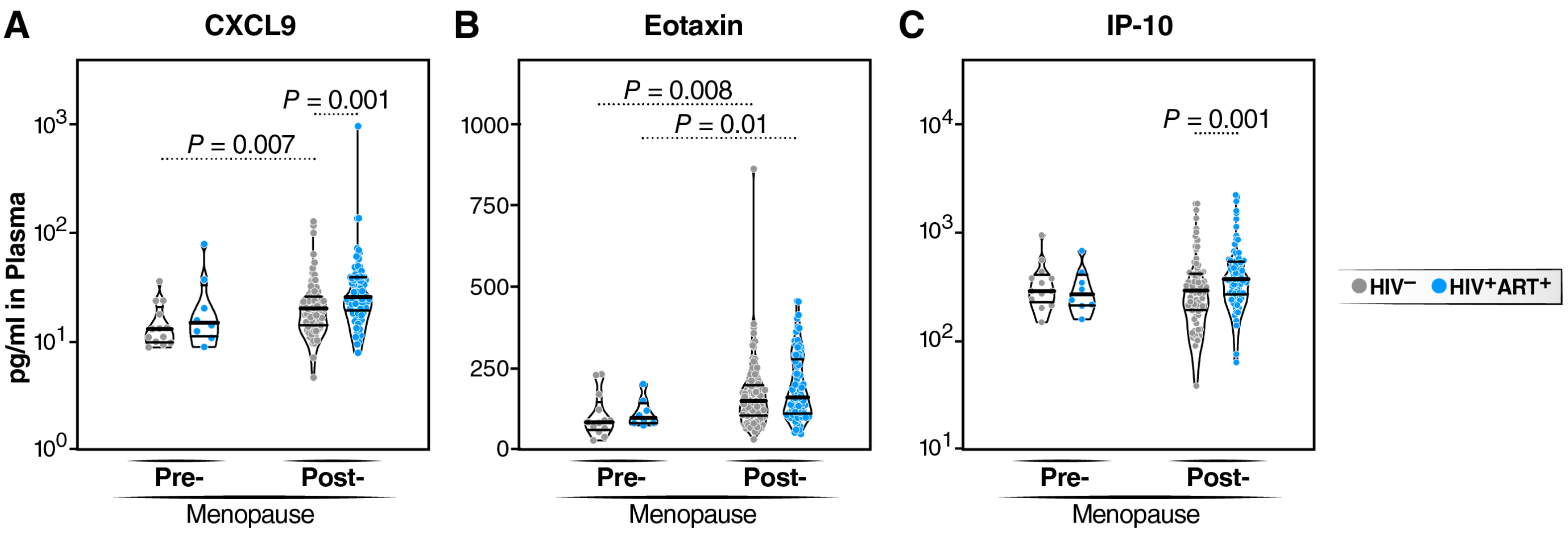

### Supplementary Figure 4

##### Supplementary Figure 4

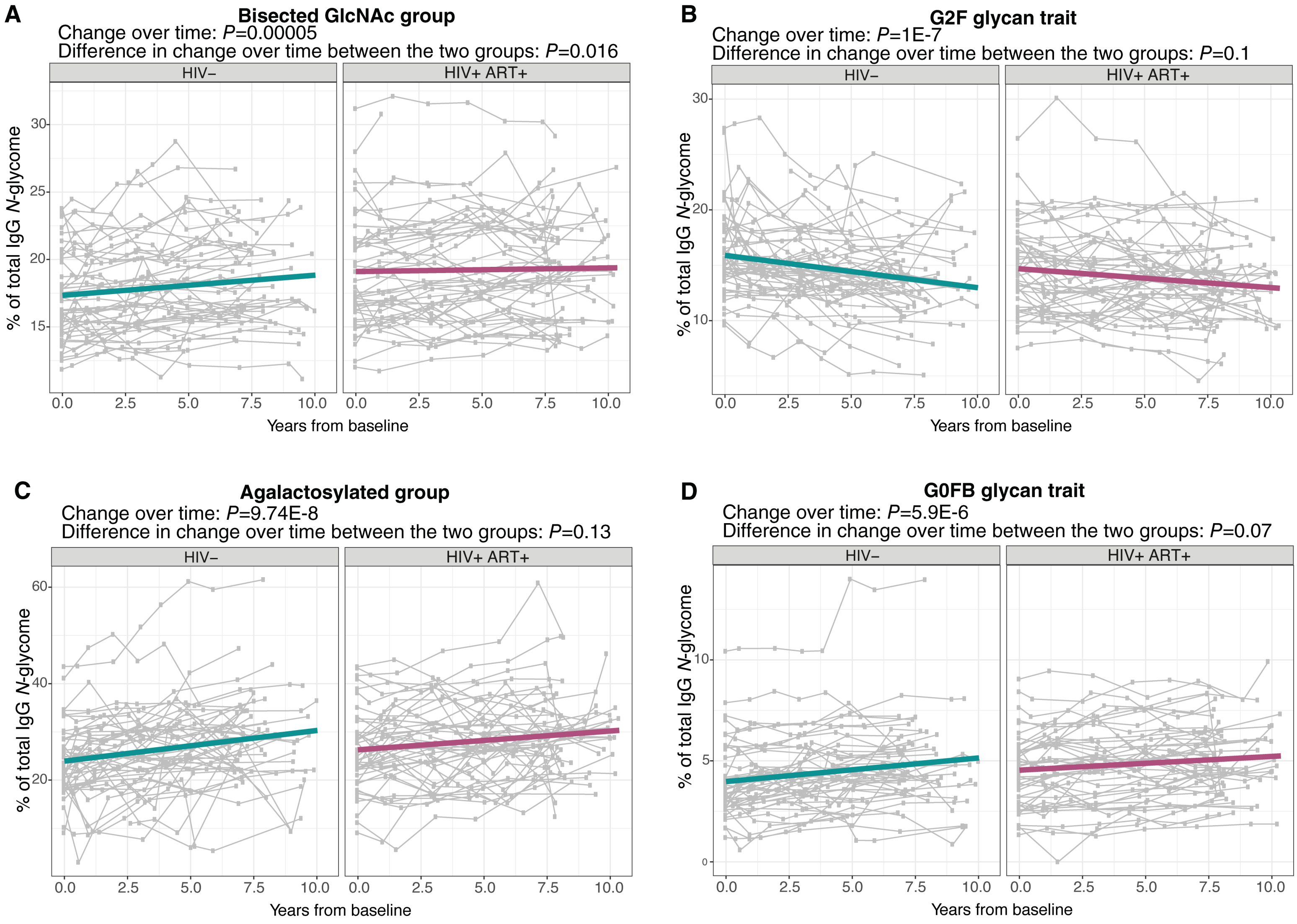

### Supplementary Figure 5

##### Supplementary Figure 5

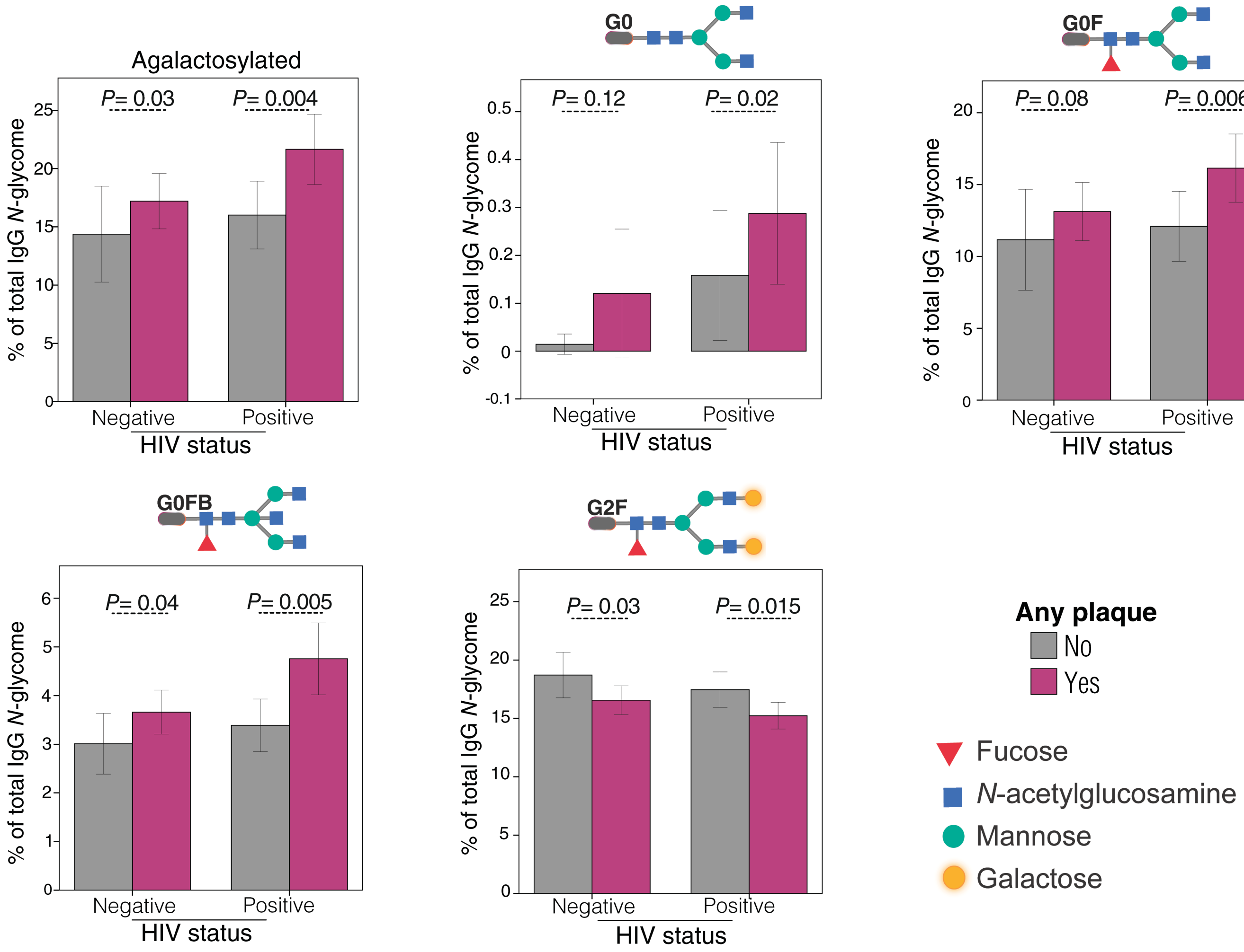

### Supplementary Figure 6

**Supplementary Figure 6**

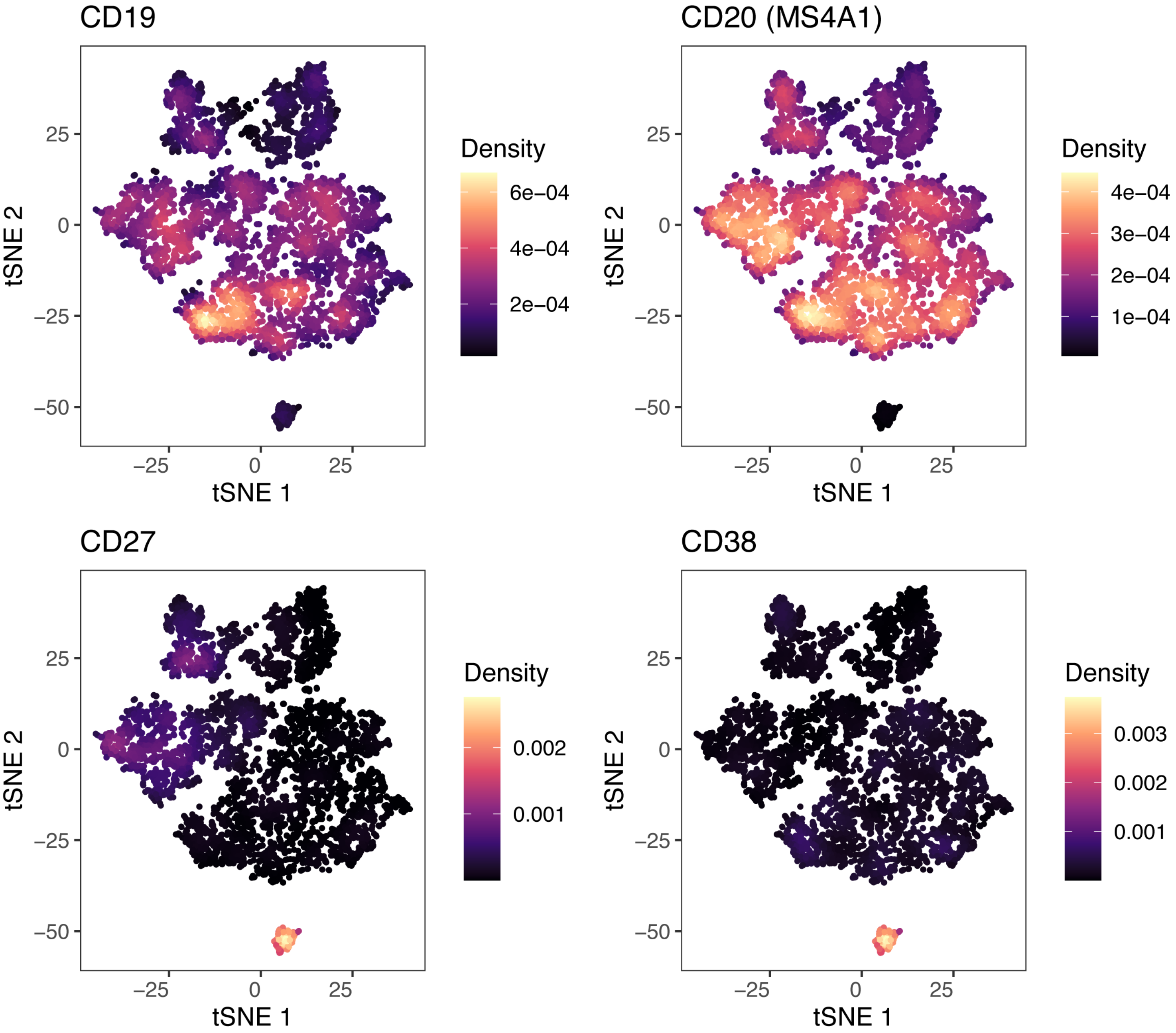

### Supplementary Figure 7

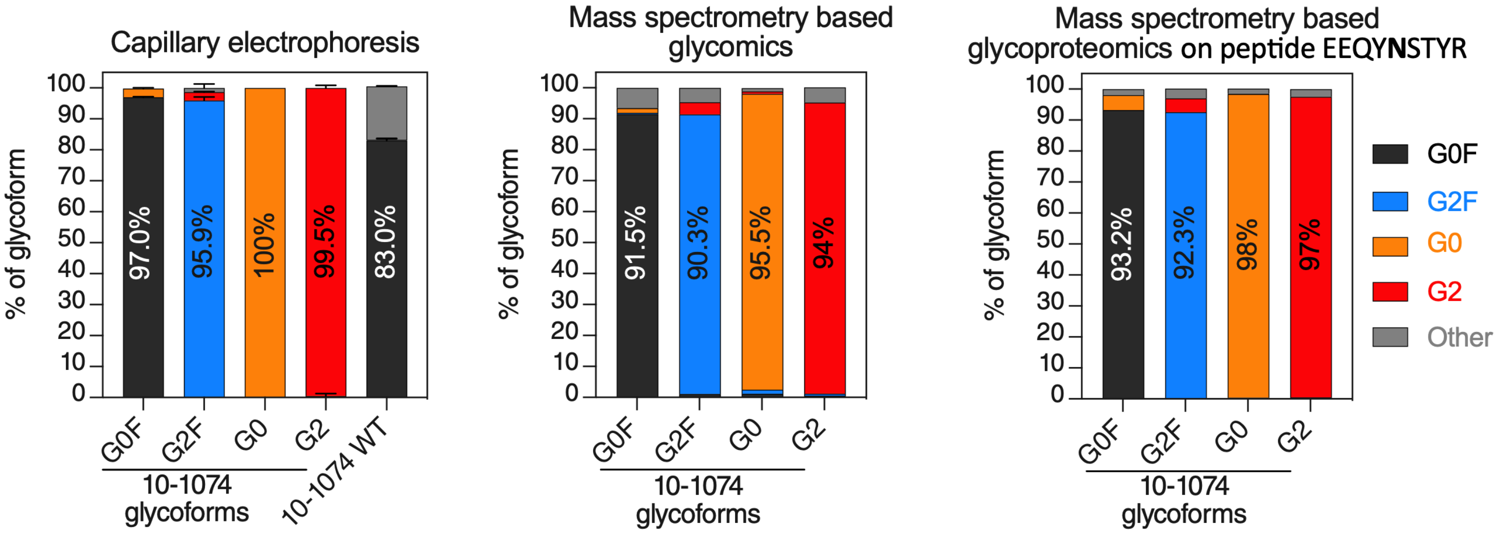
