## Supplementary Table 1 for "Plasma Glycomic Markers of Accelerated Biological Aging During Chronic HIV Infection"

**Supplementary Table 1.** Comparisons of IgG glycans among pre-menopause, peri-menopause, and post-menopause women.

| Glycan Trait | Pre-menopause | | | Early/Late peri-menopause | | | Post-menopause | | | HIV negative pre-menoapuse | | HIV+ ART+ pre-menoapuse | |
|---|---|---|---|---|---|---|---|---|---|---|---|---|---|
| | HIV- | HIV+ ART+ | | HIV- | HIV+ ART+ | | HIV- | HIV+ ART+ | | *vs* HIV$^-$ peri-menopause | *vs* *HIV$^-$* Post-menopause | *vs* HIV$^+$ peri-menopause | *vs* *HIV$^+$* Post-menopause |
| | Median | Median | *P* value | Median | Median | *P* value | Median | Median | *P* value | *P* value | *P* value | *P* value | *P* value |
| A2 | 0.94% | 0.93% | 0.55 | 0.86% | 1.20% | **0.018** | 0.88% | 0.97% | **0.002** | KW non-significant | | KW non-significant | |
| A2B | 0.18% | 0.29% | **0.002** | 0.08% | 0.29% | **0.016** | 0.22% | 0.25% | 0.054 | 0.834 | 0.200 | KW non-significant | |
| A2F | 2.15% | 2.04% | 0.305 | 1.67% | 1.97% | 0.137 | 1.52% | 1.64% | **0.04** | **0.009** | **<0.0001** | 1.000 | **0.011** |
| A2FB | 2.17% | 2.38% | 0.321 | 2.15% | 2.44% | 0.200 | 2.22% | 2.26% | 0.12 | KW non-significant | | KW non-significant | |
| A1 | 0.87% | 0.97% | 0.790 | 0.62% | 0.93% | 0.124 | 0.59% | 0.69% | 0.287 | 0.067 | **<0.0001** | 1.000 | **0.007** |
| A1p | 0.21% | 0.40% | **0.001** | 0.19% | 0.37% | **0.007** | 0.31% | 0.34% | **0.033** | 1.000 | **0.015** | KW non-significant | |
| A1F | 11.09% | 9.51% | **0.042** | 8.55% | 7.18% | 0.365 | 6.52% | 5.90% | **0.026** | **0.045** | **<0.0001** | **0.018** | **<0.0001** |
| G0 | 0.00% | 0.11% | **0.002** | 0.00% | 0.00% | 0.223 | 0.19% | 0.40% | **0.001** | 0.900 | **<0.0001** | 1.000 | **0.002** |
| A1FB | 2.04% | 1.99% | 0.436 | 1.91% | 1.92% | 0.821 | 1.95% | 1.95% | 0.89 | KW non-significant | | KW non-significant | |
| G0B | 0.00% | 0.18% | 0.578 | 0.00% | 0.31% | 0.317 | 0.39% | 0.43% | 0.6 | 1.000 | **<0.0001** | 0.800 | **0.007** |
| G0F | 17.10% | 18.54% | 0.547 | 22.98% | 24.08% | 0.529 | 26.29% | 27.09% | 0.09 | **0.025** | **<0.0001** | **0.009** | **<0.0001** |
| G1 | 0.54% | 1.09% | **<0.0001** | 0.60% | 0.75% | 0.224 | 0.66% | 0.86% | **0.006** | KW non-significant | | KW non-significant | |
| G0FB | 2.97% | 3.25% | 0.305 | 4.51% | 4.63% | 0.880 | 4.80% | 5.46% | 0.069 | **0.004** | **<0.0001** | **0.037** | **<0.0001** |
| G1B | 0.00% | 0.18% | **0.040** | 0.00% | 0.00% | 0.110 | 0.11% | 0.17% | 0.198 | 0.900 | **0.040** | KW non-significant | |
| G1B_2 | 0.00% | 0.00% | 0.548 | 0.00% | 0.00% | 0.600 | 0.00% | 0.00% | 0.647 | KW non-significant | | KW non-significant | |
| G1F | 21.67% | 21.81% | 0.975 | 21.37% | 21.84% | 0.650 | 21.95% | 21.36% | **0.004** | KW non-significant | | KW non-significant | |
| G1Fp | 8.73% | 8.10% | 0.137 | 8.92% | 8.14% | 0.160 | 9.27% | 7.87% | **<0.0001** | KW non-significant | | KW non-significant | |
| G1FB | 5.85% | 6.48% | **0.006** | 6.06% | 5.86% | 0.920 | 5.79% | 6.17% | **0.018** | KW non-significant | | KW non-significant | |
| G2F | 18.96% | 17.74% | 0.300 | 14.41% | 14.27% | 0.670 | 12.60% | 11.85% | 0.074 | **0.014** | **<0.0001** | **0.006** | **<0.0001** |
| G2FB | 1.58% | 1.62% | 0.123 | 1.40% | 1.39% | 0.400 | 1.25% | 1.22% | 0.947 | 1.000 | **<0.0001** | **0.007** | **<0.0001** |

KW = Kruskal Wallis test
