## Supplementary Table 2 for "Plasma Glycomic Markers of Accelerated Biological Aging During Chronic HIV Infection"

**Supplementary Table 2.** Correlations between IgG glycans and chronological age.

| | Variable | Group | Women | | | | Men | | | | All | | | |
|---|---|---|---|---|---|---|---|---|---|---|---|---|---|---|
| | | | Spearman's rho | *P* | Slope | Slope *P* | Spearman's rho | *P* | Slope | Slope *P* | Spearman's rho | *P* | Slope | Slope *P* |
| Individual glycans | G0 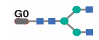 | HIV- | 0.3500 | **<0.0001** | 0.0130 | ***0.050*** | 0.4000 | **<0.0001** | 0.0140 | 0.120 | 0.3800 | **<0.0001** | 0.0140 | **0.008** |
| | | HIV+ ART+ | 0.3100 | **<0.0001** | 0.0250 | | 0.4000 | **<0.0001** | 0.0230 | | 0.3300 | **<0.0001** | 0.0240 | |
| | G1B [6] 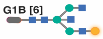 | HIV- | 0.0790 | 0.2300 | 0.0014 | 0.880 | 0.0890 | 0.1600 | 0.0009 | 0.660 | 0.0790 | 0.0800 | 0.0010 | 0.780 |
| | | HIV+ ART+ | 0.1100 | 0.0700 | 0.0013 | | 0.1840 | **0.0040** | 0.0010 | | 0.1450 | **0.0012** | 0.0012 | |
| | G1 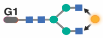 | HIV- | 0.1500 | **0.0300** | 0.0120 | 0.260 | 0.3200 | **<0.0001** | 0.0240 | 0.150 | 0.2800 | **<0.0001** | 0.0200 | 0.070 |
| | | HIV+ ART+ | -0.0011 | 0.9900 | 0.0020 | | 0.1400 | **0.0350** | 0.0120 | | 0.0900 | **0.0460** | 0.0100 | |
| | A1 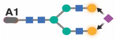 | HIV- | -0.2400 | **0.0002** | -0.0160 | 0.200 | -0.0700 | 0.2900 | -0.0100 | ***0.020*** | -0.0300 | 0.4000 | -0.0030 | ***0.045*** |
| | | HIV+ ART+ | -0.1300 | **0.0300** | -0.0090 | | -0.3000 | **0.0350** | <0.0001 | | 0.1600 | **0.0003** | -0.0180 | |
| | G1FB 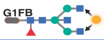 | HIV- | 0.0200 | 0.7200 | 0.0040 | 0.600 | 0.3600 | **<0.0001** | 0.0700 | ***0.030*** | 0.2200 | **<0.0001** | 0.0400 | 0.078 |
| | | HIV+ ART+ | -0.0090 | 0.8800 | -0.0060 | | 0.1200 | 0.0580 | 0.0300 | | 0.0600 | 0.1200 | 0.0200 | |
| | A2B 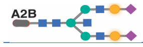 | HIV- | 0.2200 | **0.0008** | 0.0040 | ***0.017*** | 0.0400 | 0.5500 | 0.0030 | ***0.080*** | 0.2000 | **<0.0001** | 0.0060 | ***0.002*** |
| | | HIV+ ART+ | -0.0050 | 0.9400 | -0.0004 | | -0.0900 | 0.1600 | -0.0030 | | -0.0200 | 0.6400 | -0.0010 | |
| | G0FB 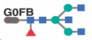 | HIV- | 0.5000 | **<0.0001** | 0.1200 | 0.710 | 0.3500 | **<0.0001** | 0.0900 | 0.400 | 0.3700 | **<0.0001** | 0.0900 | 0.690 |
| | | HIV+ ART+ | 0.4300 | **<0.0001** | 0.1300 | | 0.2700 | **<0.0001** | 0.0700 | | 0.3400 | **<0.0001** | 0.0900 | |
| | G0F 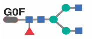 | HIV- | 0.4600 | **<0.0001** | 0.4500 | 0.760 | 0.2100 | **0.0008** | 0.1900 | 0.640 | 0.2500 | **<0.0001** | 0.2200 | 0.180 |
| | | HIV+ ART+ | 0.3800 | **<0.0001** | 0.4800 | | 0.2400 | **0.0002** | 0.2200 | | 0.2900 | **<0.0001** | 0.3000 | |
| | A2FB 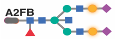 | HIV- | 0.0200 | 0.9700 | 0.0030 | 0.820 | 0.0300 | 0.6200 | 0.0040 | 0.970 | 0.0054 | 0.9000 | 0.0050 | 0.230 |
| | | HIV+ ART+ | -0.0500 | 0.4200 | -0.0010 | | 0.0620 | 0.3300 | -0.0060 | | -0.0600 | 0.1200 | -0.0040 | |
| | A2 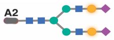 | HIV- | -0.0800 | 0.1700 | -0.0010 | 0.490 | -0.2700 | **<0.0001** | -0.0400 | 0.580 | -0.0500 | 0.2200 | -0.0050 | 0.440 |
| | | HIV+ ART+ | -0.0700 | **0.0240** | -0.0050 | | -0.1700 | **0.0060** | -0.0300 | | -0.1000 | **0.0400** | -0.0100 | |
| | G0B 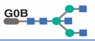 | HIV- | 0.4100 | **<0.0001** | 0.0140 | 0.800 | 0.3300 | **<0.0001** | 0.0100 | 0.380 | 0.3500 | **<0.0001** | 0.0110 | 0.260 |
| | | HIV+ ART+ | 0.2800 | **<0.0001** | 0.0130 | | 0.3200 | **<0.0001** | 0.0130 | | 0.3100 | **<0.0001** | 0.0140 | |
| | A1 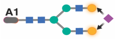 | HIV- | 0.2600 | **<0.0001** | 0.0070 | ***0.003*** | 0.1500 | **0.0200** | 0.0040 | ***0.038*** | 0.2100 | **<0.0001** | 0.0060 | ***0.001*** |
| | | HIV+ ART+ | -0.0100 | 0.8400 | -0.0010 | | -0.0100 | 0.8800 | -0.0010 | | 0.0010 | 0.9600 | -0.0004 | |
| | A1F 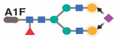 | HIV- | -0.2700 | **<0.0001** | -0.1100 | 0.420 | -0.5100 | **<0.0001** | -0.2400 | 0.110 | -0.3300 | **<0.0001** | -0.1400 | 0.440 |
| | | HIV+ ART+ | -0.2600 | **<0.0001** | -0.0800 | | -0.4500 | **<0.0001** | -0.1900 | | -0.3500 | **<0.0001** | -0.1200 | |
| | G1F [3] 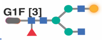 | HIV- | -0.0300 | 0.6000 | -0.0110 | 0.090 | -0.1800 | **0.0060** | -0.0600 | 0.990 | -0.1800 | **<0.0001** | -0.0640 | 0.940 |
| | | HIV+ ART+ | -0.1700 | **0.0060** | -0.0600 | | -0.1600 | **0.0100** | -0.0600 | | -0.1700 | **0.0002** | -0.0660 | |
| | G2F 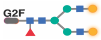 | HIV- | -0.5000 | **<0.0001** | -0.3300 | 0.770 | -0.2500 | **<0.0001** | -0.1300 | 0.630 | -0.3400 | **<0.0001** | -0.1900 | 0.460 |
| | | HIV+ ART+ | -0.4900 | **<0.0001** | -0.3200 | | -0.3400 | **<0.0001** | -0.1500 | | -0.4100 | **<0.0001** | -0.2100 | |
| | G1F [6] 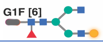 | HIV- | -0.0300 | 0.6900 | -0.0020 | 0.260 | -0.1000 | 0.1200 | -0.0200 | ***0.006*** | -0.0200 | 0.6400 | 0.0010 | 0.400 |
| | | HIV+ ART+ | 0.1400 | **0.0300** | -0.0200 | | -0.1800 | **0.0060** | 0.0300 | | 0.0400 | 0.3800 | 0.0100 | |
| | A2F 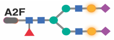 | HIV- | -0.3800 | **<0.0001** | -0.0360 | 0.400 | -0.2600 | **<0.0001** | -0.0300 | 0.310 | -0.2500 | **<0.0001** | -0.0230 | 0.870 |
| | | HIV+ ART+ | -0.3000 | **<0.0001** | -0.0300 | | -0.2500 | **<0.0001** | -0.0200 | | -0.2700 | **<0.0001** | -0.0240 | |
| | G2FB 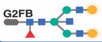 | HIV- | -0.2500 | **0.0001** | -0.0100 | 0.170 | 0.0800 | 0.1900 | -0.1100 | 0.140 | -0.0080 | 0.8600 | 0.0005 | ***0.010*** |
| | | HIV+ ART+ | -0.3100 | **<0.0001** | -0.0200 | | -0.1100 | 0.0800 | -0.0050 | | -0.1900 | **<0.0001** | -0.0100 | |
| | A1FB 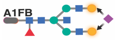 | HIV- | -0.0700 | 0.3100 | -0.0040 | 0.950 | -0.0600 | 0.3000 | -0.0070 | 0.440 | 0.0050 | 0.9000 | 0.0040 | 0.070 |
| | | HIV+ ART+ | -0.0080 | 0.2100 | -0.0030 | | -0.2100 | **0.0080** | -0.1500 | | -0.1300 | **0.0040** | -0.0080 | |
| | G1B [3] 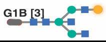 | HIV- | 0.2200 | **0.0005** | 0.0050 | 0.140 | 0.3500 | **<0.0001** | 0.0080 | 0.630 | 0.3200 | **<0.0001** | 0.0800 | 0.330 |
| | | HIV+ ART+ | 0.0500 | 0.4000 | 0.0010 | | 0.2000 | **0.0020** | 0.0070 | | 0.1500 | **0.0010** | 0.0060 | |
| Grouped glycans | **Bisected** | HIV- | 0.3000 | **<0.0001** | 0.1300 | 0.640 | 0.3400 | **<0.0001** | 0.1700 | 0.058 | 0.3500 | **<0.0001** | 0.1600 | 0.059 |
| | | HIV+ ART+ | 0.2200 | **0.0003** | 0.1100 | | 0.1800 | **0.0040** | 0.0800 | | 0.2000 | **<0.0001** | 0.1000 | |
| | **Fucosylated** | HIV- | -0.2300 | **0.0004** | -0.0540 | 0.500 | -0.0960 | 0.1270 | -0.2200 | 0.410 | -0.2500 | **<0.0001** | -0.0700 | 0.130 |
| | | HIV+ ART+ | -0.1000 | 0.0900 | -0.0300 | | 0.0100 | 0.8500 | 0.0070 | | -0.1000 | **0.0400** | -0.0300 | |
| | **Agalactosylated** | HIV- | 0.5000 | **<0.0001** | 0.6000 | 0.670 | 0.2600 | **<0.0001** | 0.3000 | 0.740 | 0.2900 | **<0.0001** | 0.3300 | 0.160 |
| | | HIV+ ART+ | 0.4300 | **<0.0001** | 0.6500 | | 0.2900 | **<0.0001** | 0.3300 | | 0.3300 | **<0.0001** | 0.4300 | |
| | **Terminal G ratio** | HIV- | 0.5000 | **<0.0001** | 0.0110 | 0.150 | 0.2600 | **<0.0001** | 0.0050 | 0.300 | 0.3000 | **<0.0001** | 0.0060 | ***0.019*** |
| | | HIV+ ART+ | 0.4300 | **<0.0001** | 0.0150 | | 0.3000 | **<0.0001** | 0.0070 | | 0.3500 | **<0.0001** | 0.0100 | |
| | **Terminal galactosylated** | HIV- | -0.5100 | **0.0006** | -0.5900 | 0.720 | -0.2200 | **<0.0001** | -0.2300 | 0.590 | -0.3000 | **<0.0001** | -0.3200 | 0.260 |
| | | HIV+ ART+ | -0.4400 | **<0.0001** | 0.6300 | | -0.2700 | **<0.0001** | -0.2700 | | -0.3400 | **<0.0001** | -0.4000 | |
| | **Sialylated** | HIV- | -0.3000 | **<0.0001** | -0.2000 | 0.960 | -0.3000 | **<0.0001** | -0.3200 | 0.350 | -0.2700 | **<0.0001** | -0.3100 | 0.380 |
| | | HIV+ ART+ | -0.3200 | **<0.0001** | -0.1900 | | -0.3200 | **<0.0001** | -0.1900 | | -0.1500 | **<0.0001** | -0.1900 | |

■ *N*-acetylglucosamine ▲ Fucose ● Mannose ● Galactose ◆ Sialic acid
