## Supplementary Table 3 for "Plasma Glycomic Markers of Accelerated Biological Aging During Chronic HIV Infection"

**Supplementary Table 3.** Characteristics of participants in the longitudinal analysis presented in Supplementary Figure 4.

| | | | Visit 1 | | Visit 2 | | Visit 3 | | Visit 4 | | Visit 5 | | Visit 6 | | Visit 7 | |
|---|---|---|---|---|---|---|---|---|---|---|---|---|---|---|---|---|
| | | | HIV- | HIV+ | HIV- | HIV+ | HIV- | HIV+ | HIV- | HIV+ | HIV- | HIV+ | HIV- | HIV+ | HIV- | HIV+ |
| **Women** | **n** | | 23 | 23 | 23 | 23 | 23 | 23 | 23 | 23 | 23 | 23 | 23 | 23 | 18 | 21 |
| | **Age (years, median IQR)** | | 49 (5.5) | 50 (5.5) | 50 (6) | 51 (5) | 51 (6) | 53 (5.5) | 52 (5.5) | 54 (6) | 54 (5.5) | 56 (5.5) | 56 (5.5) | 57 (6) | 56 (5.8) | 58 (5) |
| | **BMI (median, IQR)** | | 30 (8.2) | 29.8 (8.2) | 30 (8.7) | 39.6 (7.6) | 28 (9) | 29.8 (8.6) | 28 (9.6) | 30.1 (11.6) | 28 (8.3) | 29.9 (10) | 29 (9.2) | 29.5 (10.8) | 28.8 (9.5) | 30.3 (7.5) |
| | **CD4 Count (cells/mm3)** | | | 774 (236) | | 812 (405) | | 797 (310) | | 821 (394) | | 837 (402) | | 750 (433) | | 745 (285) |
| | **Viral Load (copies/ml)** | | - | <50 | - | <50 | - | <50 | - | <50 | - | <50 | - | <50 | - | <50 |
| | **Nadir CD4** | | - | 209 (166.5) | - | - | - | - | - | - | - | - | - | - | - | - |
| | **Race** | **White, non-Hispanic** | 4 | 4 | - | - | - | - | - | - | - | - | - | - | - | - |
| | | **Black, Hispanic** | 17 | 17 | - | - | - | - | - | - | - | - | - | - | - | - |
| | | **Other Hispanic** | 2 | 2 | - | - | - | - | - | - | - | - | - | - | - | - |
| **Men** | **n** | | 24 | 25 | 24 | 25 | 24 | 25 | 24 | 24 | 24 | 23 | 23 | 23 | 6 | 4 |
| | **Age (years, median IQR)** | | 49 (4) | 49 (4) | 50 (4) | 50 (4) | 52 (4) | 52 (4) | 54 (4) | 54 (4) | 55 (4) | 56 (5) | 57 (4) | 59 (4) | 57 (4) | 58 (1) |
| | **BMI (median, IQR)** | | 24 (4.1) | 25 (2.7) | 25 (3.7) | 25 (3.3) | 24 (4.1) | 24 (3.8) | 24 (3.8) | 25 (4.2) | 24 (3.9) | 25 (4.1) | 25 (3.8) | 25 (4.2) | 26 (2.7) | 26 (2.9) |
| | **CD4 Count (cells/mm3)** | | 882 (300) | 511 (389) | 907 (307) | 726 (354) | 958 (380) | 721 (221) | 946 (234) | 728 (222) | 897 (196) | 684 (251) | 943 (207) | 735 (379) | 911 (244) | 636 (211) |
| | **Viral Load (copies/ml)** | | - | <50 | - | <50 | - | <50 | - | <50 | - | <50 | - | <50 | - | <50 |
| | **Nadir CD4** | | 540 (138) | 277 (177) | - | - | - | - | - | - | - | - | - | - | - | - |
| | **Race** | **White, non-Hispanic** | 22 | 23 | - | - | - | - | - | - | - | - | - | - | - | - |
| | | **Black, non-Hispanic** | 2 | 2 | - | - | - | - | - | - | - | - | - | - | - | - |
