## Supplementary Table 4 for "Plasma Glycomic Markers of Accelerated Biological Aging During Chronic HIV Infection"

**Supplementary Table 4.** Correlations between inflammatory markers and chronological age.

| Inflammatory Marker | HIV- women | | | | HIV+ women | | | | HIV- men | | | | HIV+ men | | | |
|---|---|---|---|---|---|---|---|---|---|---|---|---|---|---|---|---|
| | n | Spearman's rho | *P* value | FDR | n | Spearman's rho | *P* value | FDR | n | Spearman's rho | *P* value | FDR | n | Spearman's rho | *P* value | FDR |
| **Fractalkine** | 100 | 0.225 | **0.024** | 0.106 | 100 | 0.039 | 0.698 | 0.870 | 100 | 0.131 | 0.192 | 0.534 | 100 | 0.263 | **0.008** | 0.060 |
| **IFN-α2a** | 64 | 0.059 | 0.643 | 0.884 | 50 | -0.003 | 0.982 | 0.982 | 97 | 0.108 | 0.291 | 0.640 | 94 | 0.129 | 0.217 | 0.284 |
| **IL-12p70** | 89 | 0.105 | 0.325 | 0.651 | 87 | 0.143 | 0.186 | 0.371 | 91 | 0.026 | 0.806 | 0.934 | 84 | 0.174 | 0.113 | 0.253 |
| **IL-2** | 85 | -0.006 | 0.957 | 0.957 | 73 | 0.163 | 0.168 | 0.371 | 89 | 0.139 | 0.194 | 0.534 | 86 | 0.287 | **0.007** | 0.060 |
| **IL-4** | 73 | 0.035 | 0.769 | 0.957 | 59 | -0.035 | 0.792 | 0.918 | 88 | -0.030 | 0.782 | 0.934 | 85 | 0.194 | 0.076 | 0.212 |
| **IL-5** | 92 | -0.013 | 0.899 | 0.957 | 72 | -0.044 | 0.712 | 0.870 | 92 | -0.004 | 0.967 | 0.996 | 91 | 0.157 | 0.138 | 0.253 |
| **IP-10** | 100 | 0.049 | 0.626 | 0.884 | 100 | 0.187 | 0.062 | 0.227 | 100 | 0.203 | **0.043** | 0.315 | 100 | 0.240 | **0.016** | 0.090 |
| **MCP-2** | 100 | 0.177 | 0.078 | 0.215 | 100 | 0.087 | 0.390 | 0.613 | 100 | -0.001 | 0.996 | 0.996 | 100 | -0.030 | 0.766 | 0.802 |
| **MIP-1α** | 100 | 0.068 | 0.501 | 0.884 | 98 | 0.019 | 0.851 | 0.936 | 100 | -0.061 | 0.549 | 0.805 | 99 | 0.231 | **0.022** | 0.095 |
| **SDF-1a** | 100 | 0.200 | **0.046** | 0.150 | 100 | 0.073 | 0.471 | 0.691 | 100 | 0.063 | 0.533 | 0.805 | 100 | 0.137 | 0.173 | 0.253 |
| **Eotaxin** | 100 | 0.393 | **5.26E-05** | **0.001** | 100 | 0.323 | **0.001** | **0.012** | 100 | 0.118 | 0.241 | 0.589 | 100 | -0.283 | **0.004** | 0.060 |
| **IFN-β** | 36 | 0.225 | 0.186 | 0.455 | 41 | -0.360 | **0.021** | **0.115** | 38 | -0.242 | 0.143 | 0.523 | 48 | -0.151 | 0.307 | 0.356 |
| **IFN-γ** | 93 | 0.067 | 0.525 | 0.884 | 96 | 0.098 | 0.344 | 0.583 | 100 | 0.181 | 0.071 | 0.389 | 100 | 0.116 | 0.249 | 0.304 |
| **IL-10** | 92 | 0.006 | 0.952 | 0.957 | 96 | 0.157 | 0.127 | 0.315 | 99 | 0.082 | 0.420 | 0.805 | 98 | 0.179 | 0.077 | 0.212 |
| **IL-1β** | 85 | -0.008 | 0.944 | 0.957 | 80 | 0.180 | 0.111 | 0.315 | 91 | -0.164 | 0.119 | 0.523 | 94 | 0.158 | 0.127 | 0.253 |
| **IL-21** | 22 | 0.510 | **0.015** | 0.084 | 15 | 0.309 | 0.263 | 0.482 | 65 | 0.034 | 0.789 | 0.934 | 72 | 0.165 | 0.165 | 0.253 |
| **IL-6** | 100 | 0.198 | **0.048** | 0.150 | 100 | -0.055 | 0.586 | 0.806 | 100 | -0.049 | 0.628 | 0.864 | 99 | 0.125 | 0.220 | 0.284 |
| **Leptin** | 100 | -0.049 | 0.627 | 0.884 | 98 | -0.154 | 0.129 | 0.315 | 100 | -0.075 | 0.460 | 0.805 | 100 | 0.059 | 0.560 | 0.616 |
| **CXCL9** | 100 | 0.385 | **7.58E-05** | **0.001** | 100 | 0.359 | **2.42E-04** | **0.005** | 100 | 0.342 | **0.001** | **0.011** | 100 | 0.139 | 0.168 | 0.253 |
| **TNF-α** | 100 | 0.285 | **0.004** | **0.030** | 99 | 0.207 | **0.040** | 0.175 | 100 | 0.214 | **0.033** | 0.315 | 100 | 0.158 | 0.115 | 0.253 |
| **CD14** | 100 | 0.118 | 0.240 | 0.529 | 99 | -0.011 | 0.914 | 0.957 | 100 | -0.004 | 0.966 | 0.996 | 98 | 0.023 | 0.821 | 0.821 |
| **CD163** | 100 | 0.014 | 0.892 | 0.957 | 99 | 0.289 | **0.004** | 0.027 | 100 | 0.067 | 0.506 | 0.805 | 98 | 0.200 | **0.049** | 0.179 |
