## Supplementary Table 5 for "Plasma Glycomic Markers of Accelerated Biological Aging During Chronic HIV Infection"

**Supplementary Table 5.** Baseline characteristics of the subclinical atherosclerosis study participants in studies presented in Figure 5.

| | | HIV- controls (n=22) | HIV- cases (n=22) | *P* value (HIV-, cases vs controls) | HIV+ controls (n=34) | HIV+ cases (n=34) | *P* value (HIV+, cases vs controls) |
|---|---|---|---|---|---|---|---|
| **Age (years; mean, range)** | | 57.4 (48-71) | 57.86 (49-65) | 0.654 | 51.35 (43-63) | 51.79 (44-62) | 0.653 |
| **BMI (mean, range)** | | 25.45 (20.1-33.2) | 28.66 (21-38.3) | 0.066 | 26.03 (17.7-34) | 24.94 (17.4-33.6) | 0.302 |
| **CD4 count (cells/mm³; mean, range)** | | 846.41 (288-1565) | 982.9 (551-1809) | 0.99 | 591.94 (103-1228) | 623.87 (205-1255) | 0.62 |
| **Nadir CD4 (cells/mm³; mean, range)** | | - | - | - | 282.88 (8-733) | 227.58 (0-560) | 0.19 |
| **Systolic Blood Pressure (mm Hg; mean, range)** | | 123.76 (103-148) | 130.71 (110-151) | 0.057 | 126.42 (108-164) | 124.09 (100-148) | 0.472 |
| **Diastolic Blood Pressure (mm Hg; mean, range)** | | 75.89 (62-95) | 81.95 (66-108) | **0.041** | 79.63 (65-91) | 76.45 (61-101) | 0.069 |
| **Fasting glucose (mg/dL; mean, range)** | | 93.45 (68-133) | 113 (72-247) | **0.021** | 104.6 (78-277) | 106.93 (80-252) | 0.626 |
| **Total cholesterol (mg/dL; mean, range)** | | 185.86 (138-259) | 202.31 (133-320) | 0.265 | 190.23 (115-294) | 186.63 (127-269) | 0.749 |
| **High Density Lipoprotein (mg/dL; mean, range)** | | 55.99 (38.5-90.8) | 49.44 (29-73.7) | 0.213 | 49.82 (22.9-101) | 45.18 (25-82.6) | 0.35 |
| **Triglycerides (mg/dL; mean, range)** | | 88.77 (23-199) | 168.81 (45-344) | **<0.0001** | 143.6 (43-425) | 141.36 (47-304) | 0.958 |
| **Low Density Lipoprotein (mg/dL; mean, range)** | | 112.18 (56-166) | 119.09 (52-210) | 0.639 | 113.87 (22-218) | 113.72 (65-202) | 0.818 |
| **Framingham coronary heart disease 10 yr risk (%; mean, range)** | | 7.36 (2-18) | 13.45 (3-27) | **0.004** | 8.44 (3-47) | 9.14 (2-33) | 0.27 |
| **Framingham hard coronary heart disease, 10 yr risk (%; mean,** | | 6.95 (1-12) | 13 (3-30) | **0.006** | 8 (1-30) | 8.11 (1-25) | 0.771 |
| **ACC/AHA risk estimate (mean, range)** | | 0.08 (0.027-0.225) | 0.12 (0.027-0.218) | **0.014** | 0.069 (0.012-0.291) | 0.077 (0.007-0.339) | 0.5 |
| **Cumulative years of ART use (mean, range)** | | - | - | - | 7.91 (0-15.41) | 9.3 (0.05-14.25) | 0.14 |
| **Viral suppression (%)** | **Supressed** | - | - | - | 91% | 82% | 0.48 |
| | **Non-supressed** | - | - | | 9% | 18% | |
| **Diagnosed with AIDS (%)** | **No AIDS Dx** | - | - | - | 85% | 82% | >0.9 |
| | **AIDS Dx** | - | - | | 15% | 18% | |
| **Statin use (%)** | **No** | 68% | 45% | 0.22 | 62% | 62% | >0.9 |
| | **Yes** | 32% | 55% | | 38% | 38% | |
| **Aspirin use (%)** | **No** | 27% | 27% | >0.9 | 59% | 59% | >0.9 |
| | **Yes** | 73% | 73% | | 41% | 41% | |
| **Diabetes status (%)** | **No** | 95% | 77% | 0.18 | 79% | 74% | >0.9 |
| | **Yes** | 5% | 23% | | 18% | 21% | |
| | **Insufficient data** | 0% | 0% | | 3% | 6% | |
| **Cancer diagnosis (%)** | **No Cancer Dx** | 95% | 82% | 0.34 | 82% | 82% | >0.9 |
| | **Cancer Dx** | 5% | 18% | | 18% | 18% | |
| **Smoking Status At Visit (%)** | **Never smoked** | 36% | 23% | - | 50% | 18% | - |
| | **Former smoker** | 59% | 59% | | 32% | 53% | |
| | **Current smoker** | 5% | 18% | | 18% | 24% | |
| **Race (%)** | **White, non-Hispanic** | 82% | 82% | - | 62% | 62% | - |
| | **White, Hispanic** | 5% | 5% | | 3% | 3% | |
| | **Black, non-Hispanic** | 14% | 14% | | 21% | 21% | |
| | **Black, Hispanic** | 0% | 0% | | 9% | 0% | |
| | **Other** | 0% | 0% | | 3% | 0% | |
| | **Other Hispanic** | 0% | 0% | | 3% | 15% | |
