## Supplementary Table 6 for "Plasma Glycomic Markers of Accelerated Biological Aging During Chronic HIV Infection"

**Supplementary Table 6.** Plasma markers of inflammation in of the subclinical atherosclerosis study participants in studies presented in Figure 5.

| | **HIV- controls (n=22)** | **HIV- cases (n=22)** | **HIV+ controls (n=34)** | **HIV+ cases (n=34)** | **ANOVA/ KW Test** |
|---|---|---|---|---|---|
| | Mean (SD) pg/ml | Mean (SD) pg/ml | Mean (SD) pg/ml | Mean (SD) pg/ml | *P-* value |
| **sCD163** | 548 (188.9) | 600.3 (213.7) | 683.3 (391.7) | 883.9 (630.3) | **0.020** |
| **sCD14** | 1633137 (367471) | 1580028 (348586) | 1900147 (549256) | 2011467 (470251) | **0.001** |
| **Fractalkine** | 5929.8 (1559.5) | 5374.9 (1025.3) | 6811.5 (2388.3) | 6273.8 (1758.3) | **0.034** |
| **IL10** | 0.25 (0.24) | 0.53 (0.89) | 0.33 (0.35) | 0.43 (0.57) | 0.447 |
| **IL1β** | 0.81 (2) | 0.29 (0.53) | 0.2 (0.15) | 0.24 (0.39) | 0.996 |
| **IL6** | 0.76 (0.33) | 1.1 (0.95) | 1.27 (1.941) | 1.59 (2.67) | 0.435 |
| **MCP2** | 21.04 (7.38) | 22.86 (5.96) | 28.09 (13.27) | 30.43 (21.94) | **0.007** |
| **MIP1α** | 18 (10.31) | 17.89 (6.343) | 18.37 (10.38) | 46.08 (133.72) | 0.908 |
| **SDF1α** | 734.5 (415.4) | 846.1 (545.8) | 761.5 (443.1) | 839.1 (254.2) | 0.216 |
| **TNFα** | 1.7 (1.6) | 1.53 (0.79) | 1.73 (1.3) | 2.68 (3.63) | 0.632 |
| **TGFβ1** | 7397.7 (6398.9) | 6691.3 (6832.1) | 7299.5 (8438.1) | 9173.2 (10822.5) | 0.490 |
| **TGFβ2** | 36.84 (25.2) | 37.51 (29.7) | 45.27 (54.9) | 66.64 (82.3) | 0.124 |
| **TGFβ3** | 2.73 (1.47) | 2.86 (1.55) | 3.06 (2.56) | 3.092 (2.35) | 0.810 |
