## Supplementary Table 7 for "Plasma Glycomic Markers of Accelerated Biological Aging During Chronic HIV Infection"

**Supplementary Table 7.** Characteristics of participants in studies presented in Figure 6.

| | | Cases | | Control | | *P* value |
|---|---|---|---|---|---|---|
| | | n (%) | Median, IQR | n (%) | Median, IQR | |
| **n** | | 10 | | 13 | | |
| **Race** | Black/African-American | 5 (50) | - | 6 (46.2) | - | |
| | Not Hispanic or Latino | 1 (10) | - | 1 (7.6) | - | |
| | White/Caucasian | 4 (40) | - | 6 (46.2) | - | |
| **CD4 Count (cells/mm3)** | | - | 497 (334) | - | 873 (328) | **0.0025** |
| **Viral Load (HIV copies/ml)** | | - | < 50 | - | < 50 | |
| **Nadir CD4 (cells/mm$^3$)** | | - | 153 (158) | - | 531 (290) | **0.0002** |
